# A method for filtering abnormal modified base calling in Oxford Nanopore Technologies sequencing

**DOI:** 10.1101/2024.05.07.593054

**Authors:** Ruikai Jia, Yusha Wang, Hua Ye, Xiaoshu Ma

## Abstract

**Background:** gene synthesis sequencing using the long-read Oxford Nanopore Technologies (ONT) provides a cost-effective option for gene synthesis quality control. Despite the advantage of using long reads, however, accurate base calling is influenced by modified bases.

**Results:** We introduce a method for filtering abnormal modified base calling in Oxford Nanopore Technologies sequencing. This method is based on the mapping results and perform an exact binomial test on the proportion of single base forward and reverse chain depth to determine the presence of abnormal modified base calling. Based on Sanger sequencing results, this method detected 96.79% of effective abnormal modified base calling, and the accuracy of gene synthesis sequencing increased from 99.40% before calibration to 99.98% after calibration, significantly reducing the proportion of requiring secondary gene synthesis sequencing.

**Conclusion:** This method can accurately filter abnormally modified base calling sites, reduce the proportion of using Sanger for secondary sequencing, thereby saving costs, and improving efficiency.

## Introduction

Gene synthesis is a technology that combines chemically synthesized oligo nucleotides ^[1]^ into gene fragment lengths through certain methods ^[2]^. After gene synthesis is completed, sequencing confirmation of the synthesized sequence is also required. One of the criteria for determining successful gene synthesis is whether the Sanger ^[3]^ sequencing results are 100% consistent with the target gene. Sanger is the gold standard for ultra-high precision sequencing, but there are issues with high cost and low throughput.

Compared to Sanger technology, Oxford Nanopore Technologies ^[4]^ instruments are smaller, more portable, cost-effective, and the process of constructing sequencing libraries is simpler. These advantages enable Nanopore technology to be applied to fast and real-time DNA sequencing. At the same time, Oxford Nanopore Technologies has an ultra-long sequencing read length (over 100kbp), which can be used to determine long repetitive fragments that NGS is powerless to detect. The ultra long read length of this technology makes it relatively easy to detect abnormalities such as gene insertion and deletion on the sequence.

Modified base refers to the formation of nucleotide molecules with functional groups on DNA molecules under the action of enzymes and other factors. Research has shown that modified base and unmodified identical base generate different electrical signal characteristics when passing through nanopores ^[5,6]^, which can affects the results of base calling. In this paper, we introduce a method for filtering abnormal modified base calling in Oxford Nanopore Technologies sequencing.

## Method

### DNA extraction and library construction

The plasmid DNA has been extracted and a sequencing library was generated. Library preparation and library quality control for the Nanopore platform: Fixed quantities of plasmid DNA were extracted, followed by fragmentation using transposase enzymes with integrated barcodes. Subsequently, the samples were combined and subjected to purification. The purified products were then ligated with a standardized adapter, facilitating the construction of a library compatible with the ONT sequencing platform. Once the library was appropriately diluted, it was loaded onto a sequencing chip and subjected to sequencing using the ONT platform.

### Sequencing data processing

We used Oxford Nanopore Technologies sequencing to obtain the raw electrical signal of plasmid DNA, which was stored in Fast5 format. Electrical signal data in Fast5 format is interpreted using Dorado Basecall Service software in High accuracy mode to obtain the original base information, which is stored in FastQ format. Perform quality filtering on FastQ format data, including filtering joint sequences, filtering short fragments with sequence length less than 500bp, filtering low-quality sequences with sequence quality values less than 7, and obtaining clean data after filtering.

Clean data uses minimap2 software ^[7]^ to map to the reference sequence and calculates the coverage information of sequencing data on the reference sequence, including the depth of forward and reverse chain for four types of bases at each base site. For base sites with sequencing depth of 100X or more, if the ratio of single base support to sequencing depth is less than 10%, it is considered as sequencing background noise and will not be included in subsequent analysis.

### Modified base detection

If the ratio of single base support to sequencing depth is less than 10%, it will be considered as sequencing background noise and will not be included in subsequent analysis. Using the binom.test function of the R software stats package, perform an exact binomial test on the proportion of forward and reverse chain depth for single base with sequencing depth of 10% or more. When the P-Value is less than or equal to 10^-10^, the base is judged as abnormal modified base calling.

### Evaluation methods

For the same plasmid DNA, we designed primers covering the full length based on the reference sequence and performed Sanger sequencing as a criterion for evaluating the efficacy. In this paper, the proportion of sites consistent with Sanger sequencing is used as an indicator to measure the accuracy of ONT sequencing. The sites that are inconsistent with Sanger sequencing results are used as the set that needs to be optimized to evaluate the performance of this method.

## Results

### Base calling and quality control

We used Nanopore and Sanger Technologies to sequence 4 plasmid samples. To translate raw signal data into nucleotide sequences, we conducted the base calling step using Dorado Basecall Service software(7.2.13+fba8e8925). Then we used dorado(0.5.2+7969fab) to demultiplex sequencing data for downstream analysis, and used NanoFilt (2.8.0) ^[8]^ for data visualization and processing, to assess the read-length and base calling quality. The average number of reads for each sample of Nanopore sequencing data after quality control was 2,929 ± 451.Together, the four samples median read lengths ranging from 4,016 to 8,040 bp (Fig. 1A), and median read quality ranging from 15 to 15.4 (Fig. 1B).

**Fig. 1.**
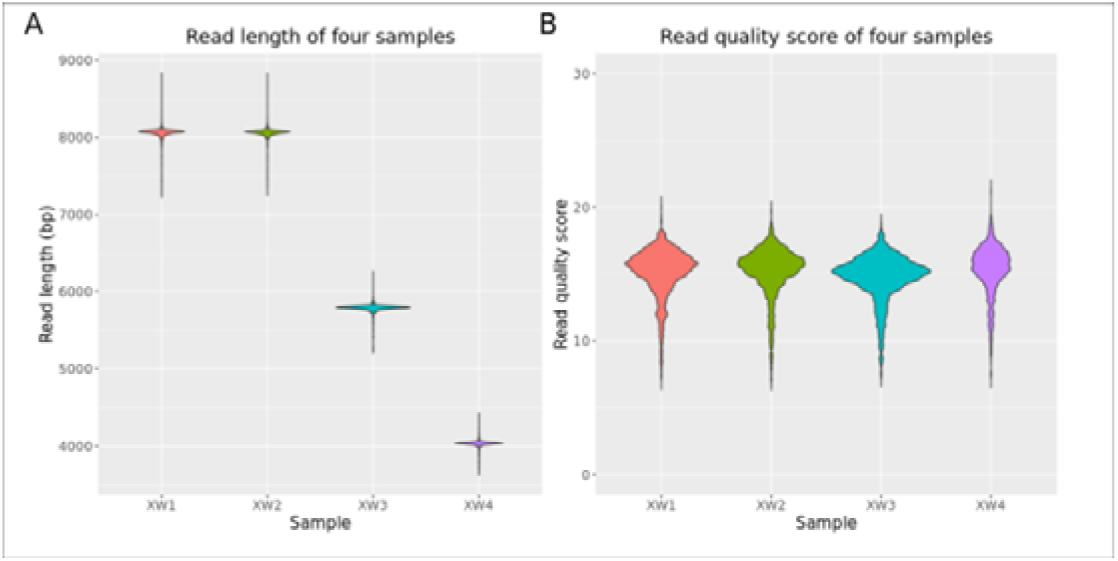
Characteristics of the nanopore sequencing. **A** Violin plot of read length near the peak. The length of sequencing data is mostly equivalent to the full length of the plasmid. Selecting data from this region to draw a Violin. **B** Violin plot of read quality score. The mean error probability (Pe) of the base was converted into a quality score using the formula: Qphred = −10log_10_Pe. Data shown are colored by Sample and plotted by R software ggplot2 package ^[9]^.

### Mapping and Modified base detection

We used Minimap2 software to map reads to reference sequences and visualized sequencing depth using Circos software (v0.69-8) ^[10]^. The sequencing depth of each site in the four samples is between 870X-2,645X (Fig. 2A), If the ratio of single base support to sequencing depth is less than 10%, it will be considered as sequencing background noise and not included in subsequent analysis. When the ratio of single base support to sequencing depth is not less than 10% and does not match the base type of the reference sequence, it is defined as a mutation candidate site (Fig. 2B). We use the binom.test function of the R software stats package to perform an exact binomial test on the proportion of forward and reverse chain depth with single bases which sequencing depth of 10% or more (Fig. 2C). Retain base types with P values greater than 10^-10^(-log10Pvalue <10), and those that are inconsistent with the reference sequence base are defined as mutation sites, as the result of single base level mutation analysis (Fig. 2D). The mutation sites also require the use of Sanger for secondary sequencing.

**Fig. 2.**
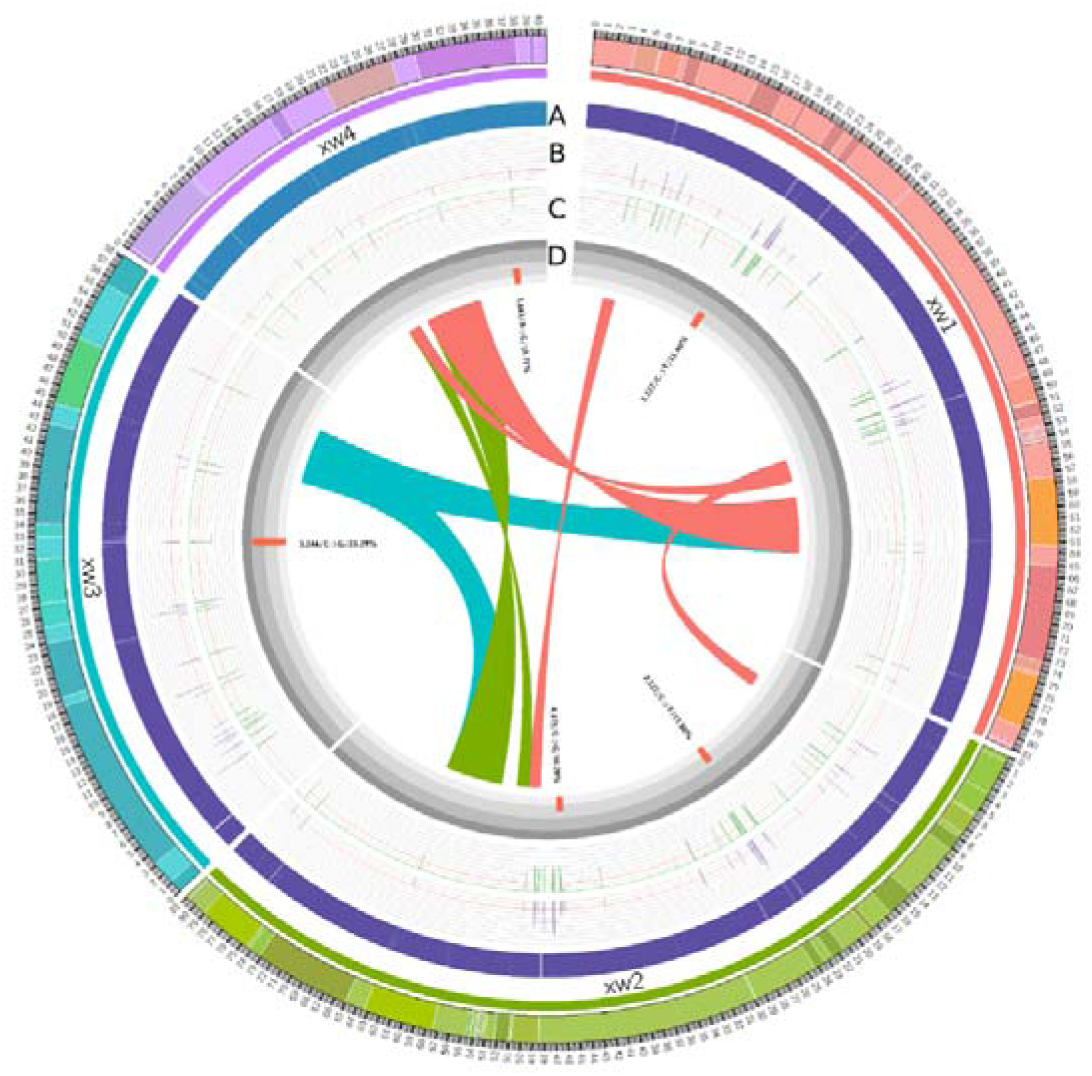
The results of Oxford Nanopore Technologies sequencing analysis. **A** Heatmap of depth distribution of sequencing data. **B** The distribution of mutation candidate sites on the reference sequence. The upper limit of the vertical axis is 50%, and the red threshold line corresponds to 10%. **C** The distribution of -log10Pvalue greater than 10 in exact binomial test results on the reference sequence. The upper limit of the vertical axis is 50, and the red threshold line is 10. **D** Distribution of mutation sites detected by Oxford Nanopore Technologies sequencing. The upper limit of the vertical axis is 30, and the displayed positions have been widened.

### Evaluation

The Sange sequencing results indicate that the plasmid DNA sample is completely consistent with the reference sequence (Supplementary Fig. S1). Among the 26,104 base sites covered by Oxford Nanopore Technologies sequencing data, there are 156 candidate polymorphic sites, and after filtering, 5 polymorphic sites remained. This indicates that the accuracy of gene synthesis sequencing using Oxford Nanopore Technologies has increased from 99.40% to 99.98%, and the filtered 151 (96.79%) candidate mutation sites can be used as a measure of the efficiency of this method. It will greatly reduce the proportion of using Sanger for secondary sequencing.

## Discussion

We proposed a method for filtering abnormal modified base calls in Oxford Nanopore Technologies sequencing data. It uses exact binomial test to determine the presence of abnormal modified base calling in sequencing data, assuming that the ratio of positive and negative chain sequences approaches 1:1 as sequencing depth increase. Due to the small size of plasmid and the requirement for high accuracy, we conducted high depth sequencing. The applicability of this method in low depth sequencing will need to be verified in the future. At the same time, we will attempt to explore the causes of abnormal modified base calling in future work.

## Conclusion

We classify the base calling in Oxford Nanopore Technologies sequencing data into three types, with the definition of abnormal modified base calling that exists due to base modification. Normal base calling is defined as one that is not at the same site as the abnormal modified base calling and is consistent with the Sanger results. Consistent with Sanger’s results, but with the presence of abnormal modified base calling at the base site, the definition of being affected by abnormal modified base calling is deviation base calling.

The ratio of sequencing depth between forward and reverse chain of normal base calling conforms to a binomial distribution (Fig. 3A), and the exact binomial test results are mostly between 0-1 after being transformed into - Log10Pvalue (Fig. 3B). The significant difference in the exact binomial test (Fig. 3C) among the three base calling types is the basis for us to use them as indicators for filtering abnormal modified base calling. For sites affected by abnormal modified base calling, receiver operating characteristic (ROC) analysis was performed using the pROC package of R software (Fig. 3D). The AUC value was 0.989, indicating that this method has extremely high accuracy.

**Fig. 3.**
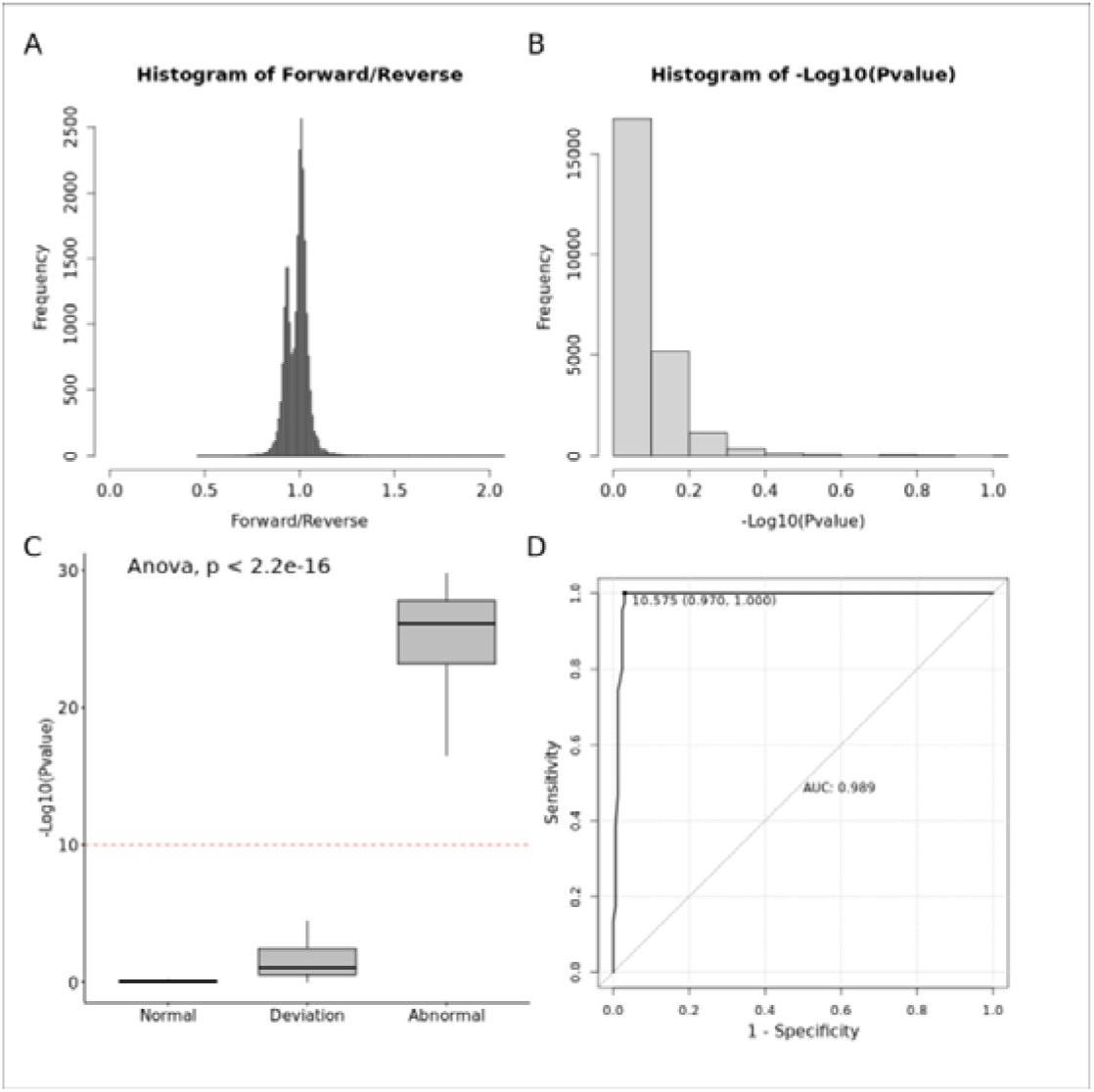
Characteristic indicators and evaluation of three types of base calling. **A** Distribution of forward/reverse of normal base calling data. The horizontal axis is the ratio of depth between forward and reverse, and the vertical axis is the frequency. **B** Distribution of exact binomial test for normal base calling. The horizontal axis represents the P value after -log10 conversion, and the vertical axis represents the frequency. **C** Comparison of three types of base calling exact binomial test results. **D** ROC analysis of sites affected by abnormal modified base calling.

In this study, we provide a method based on Oxford Nanopore Technologies sequencing data to filter out abnormal modified base calling, reducing the proportion of secondary sequencing using Sanger sequencing and improving sequencing efficiency.

## ACKNOWLEDGMENTS

We thank members of Azenta for their technical support in genome sequencing.

## Supplement

Fig. S1: Sanger result

## Data and resource sharing and availability statement

The raw sequencing data that support the findings of this study is available from China National GeneBank Sequence Archieve (CNSA) under the accession number of CNP0005663.

---

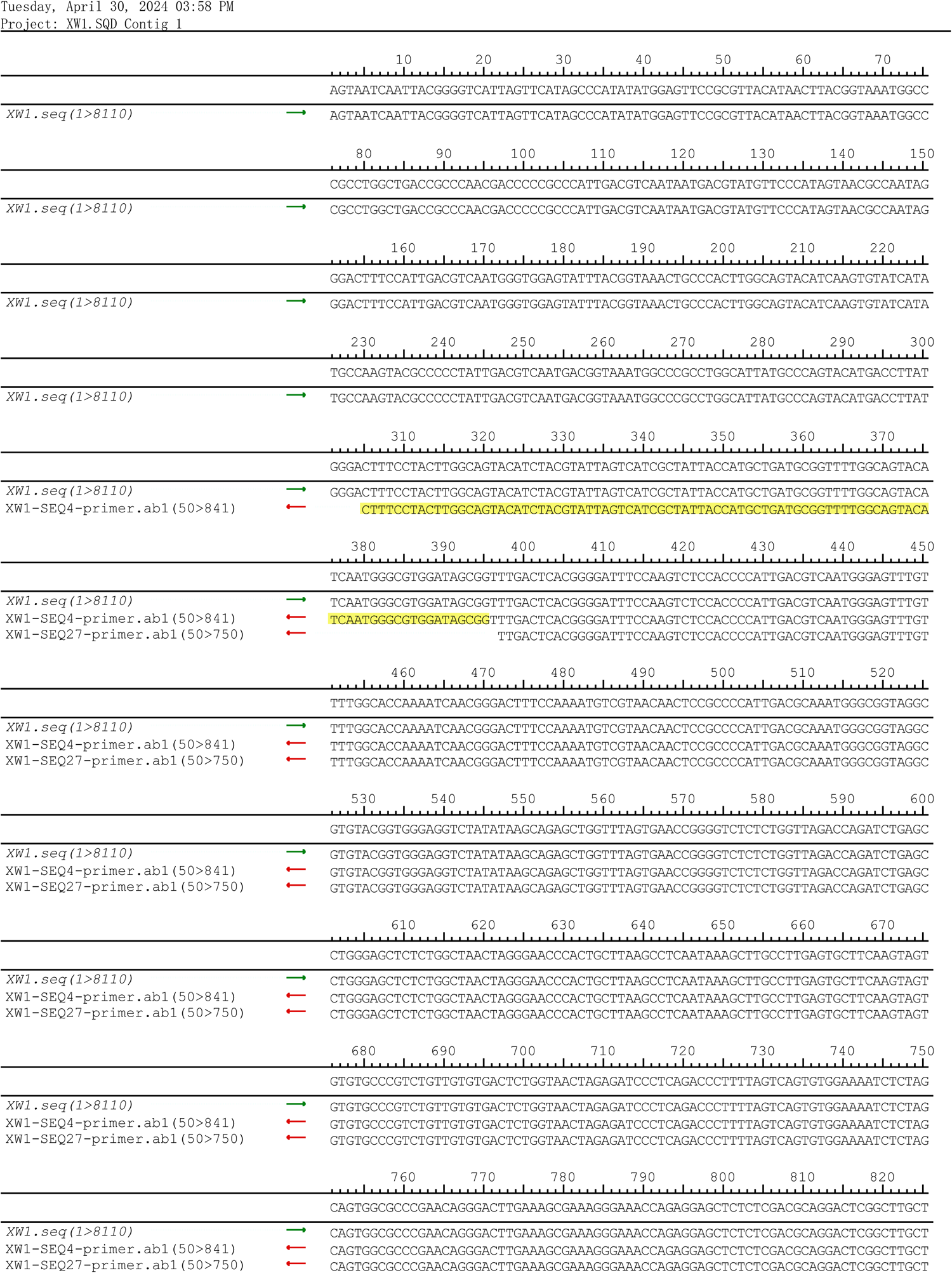

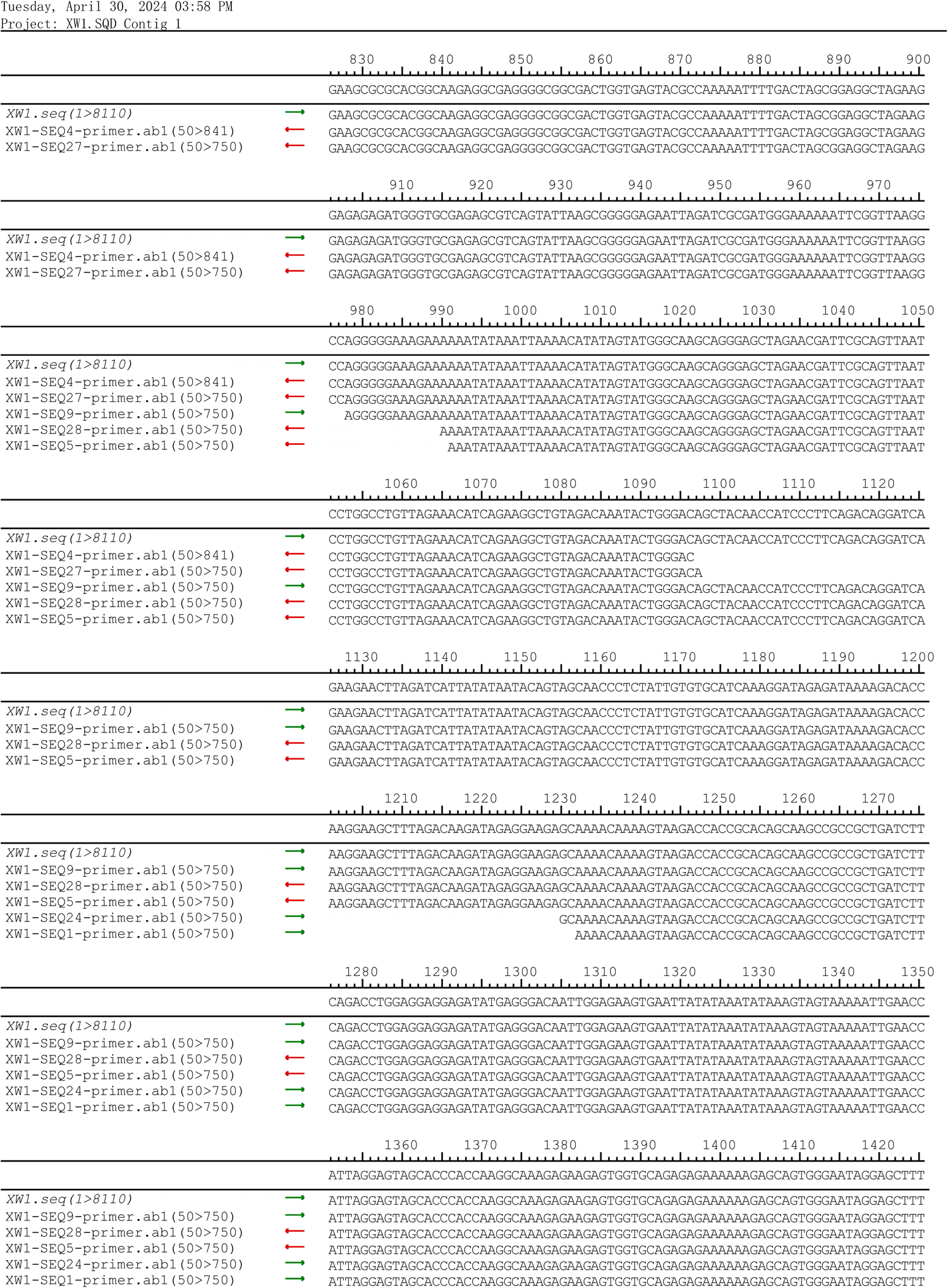

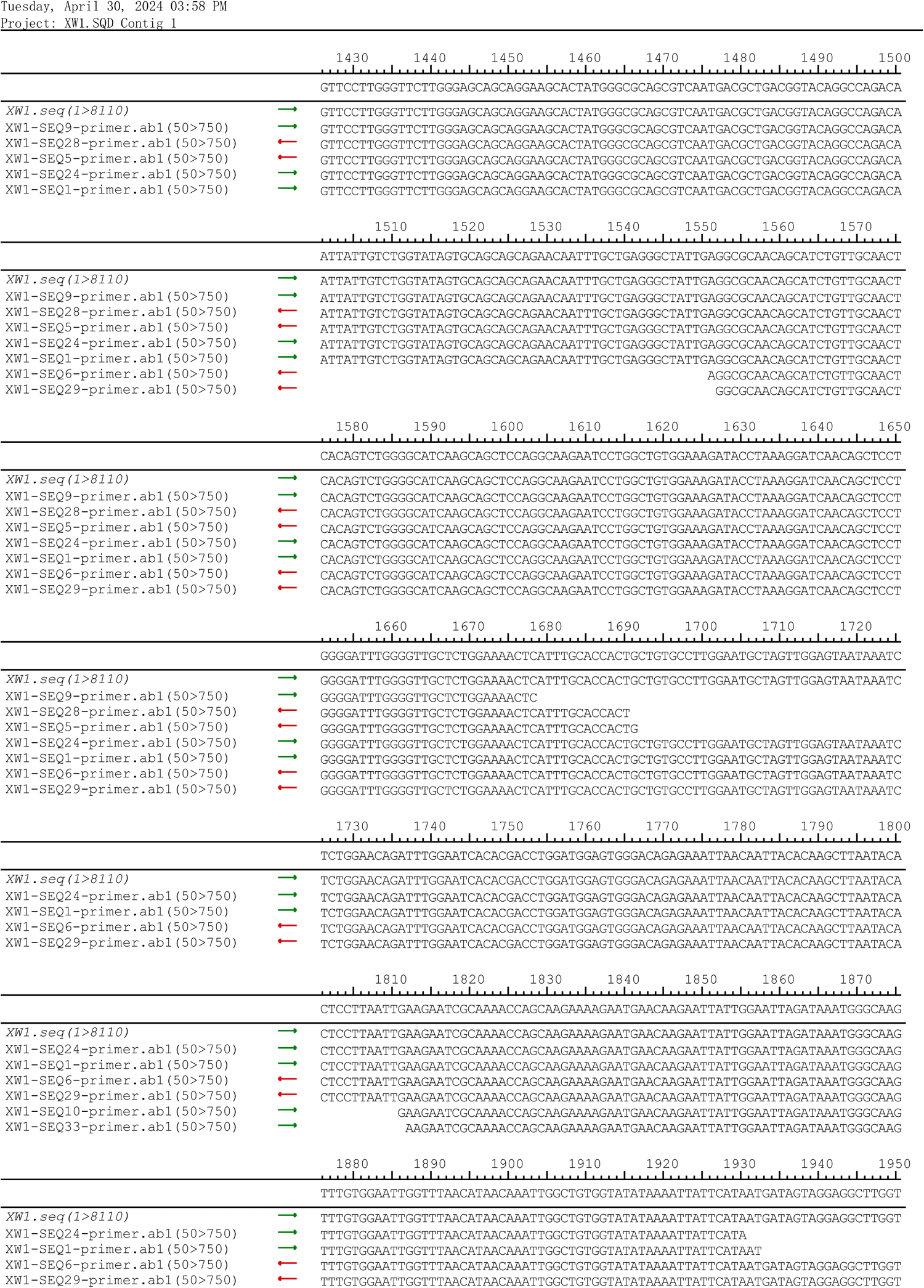

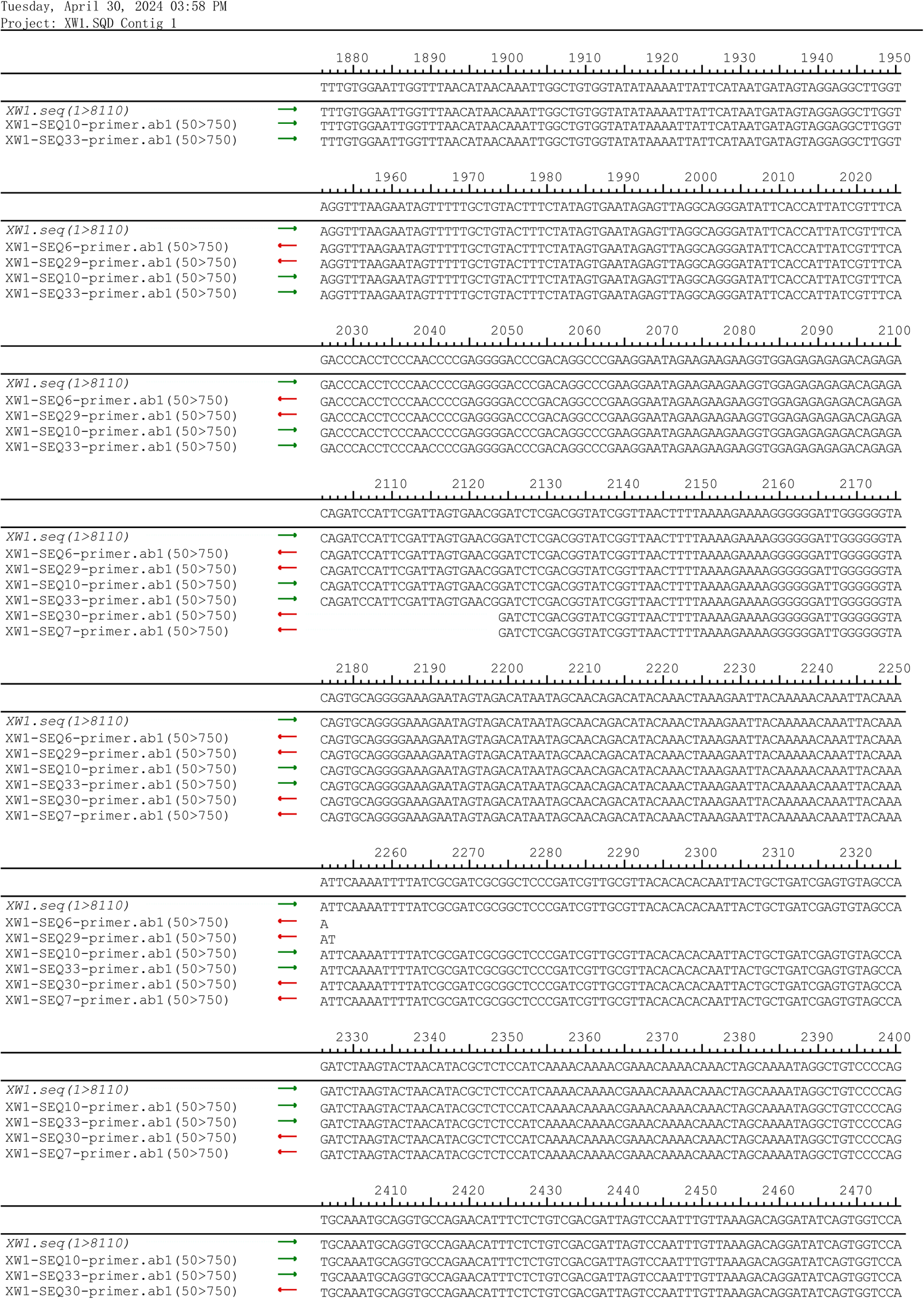

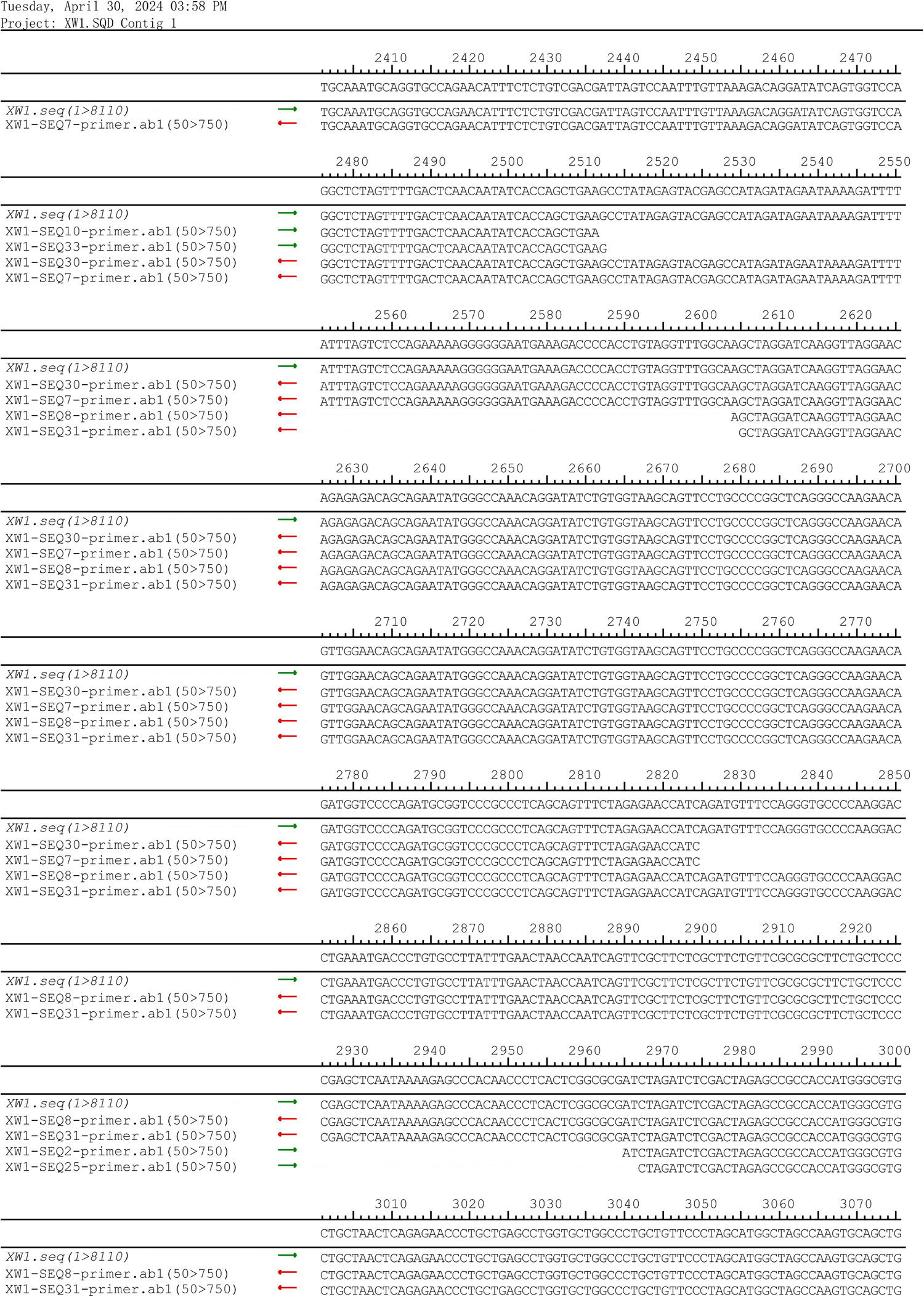

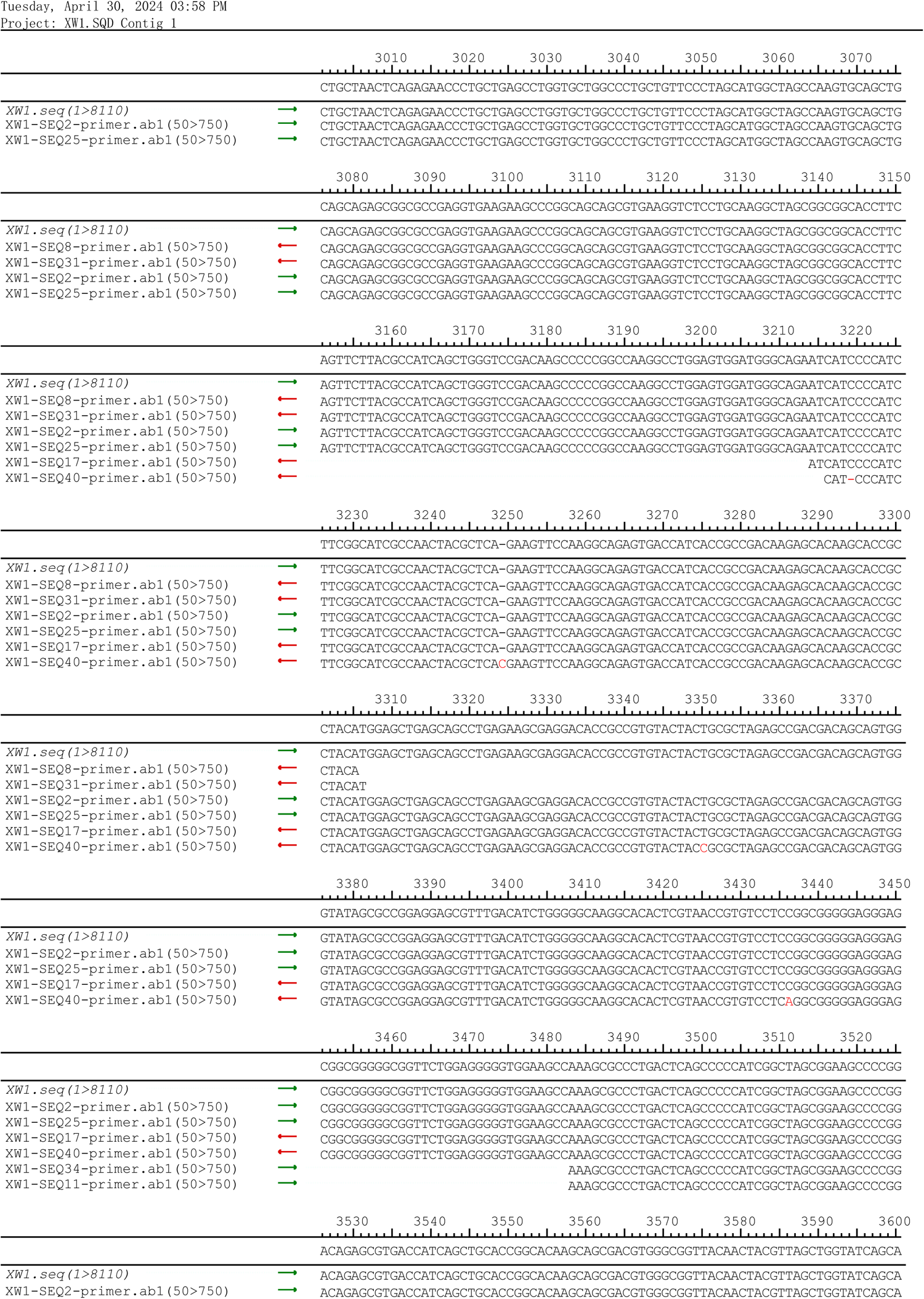

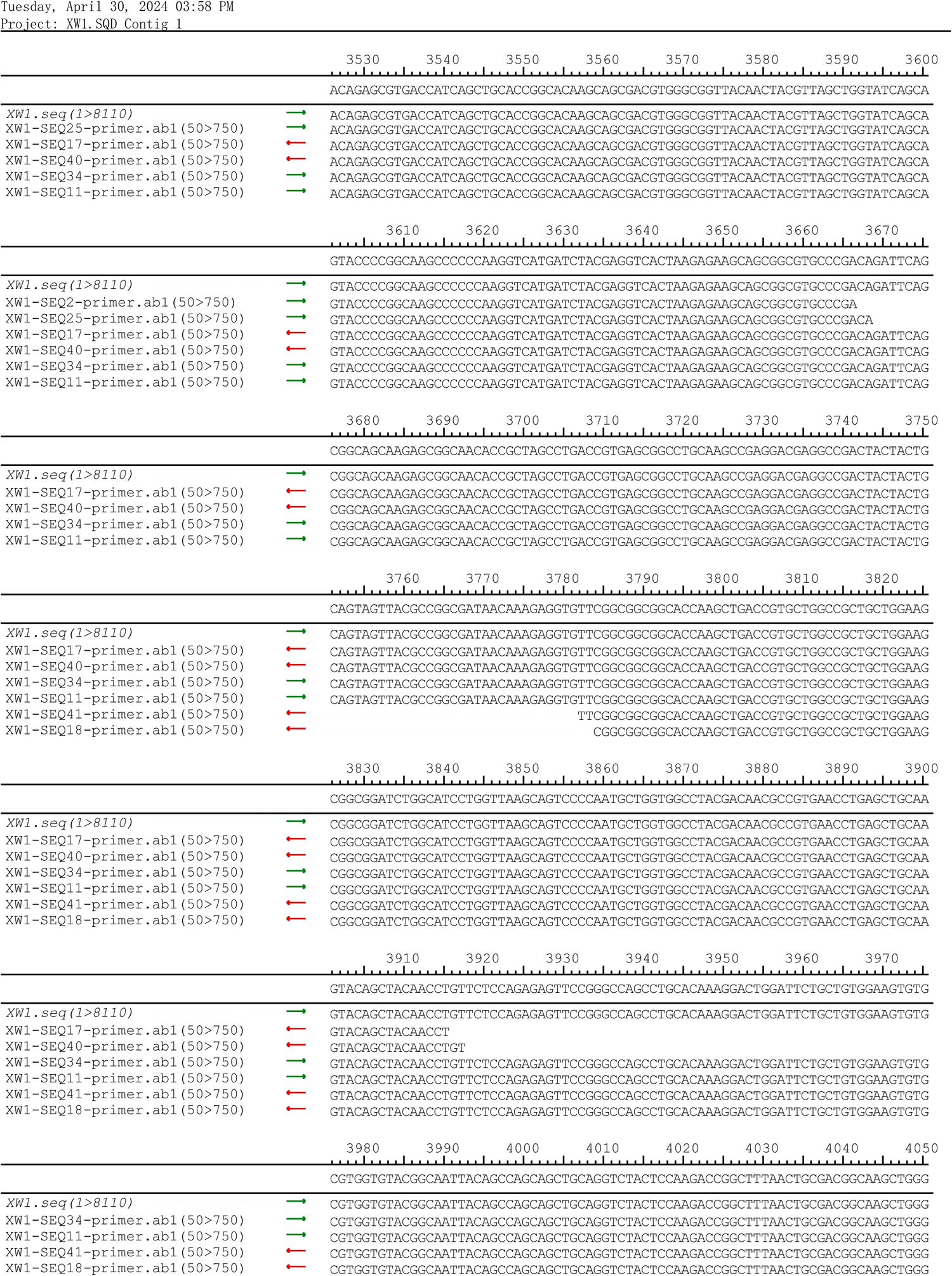

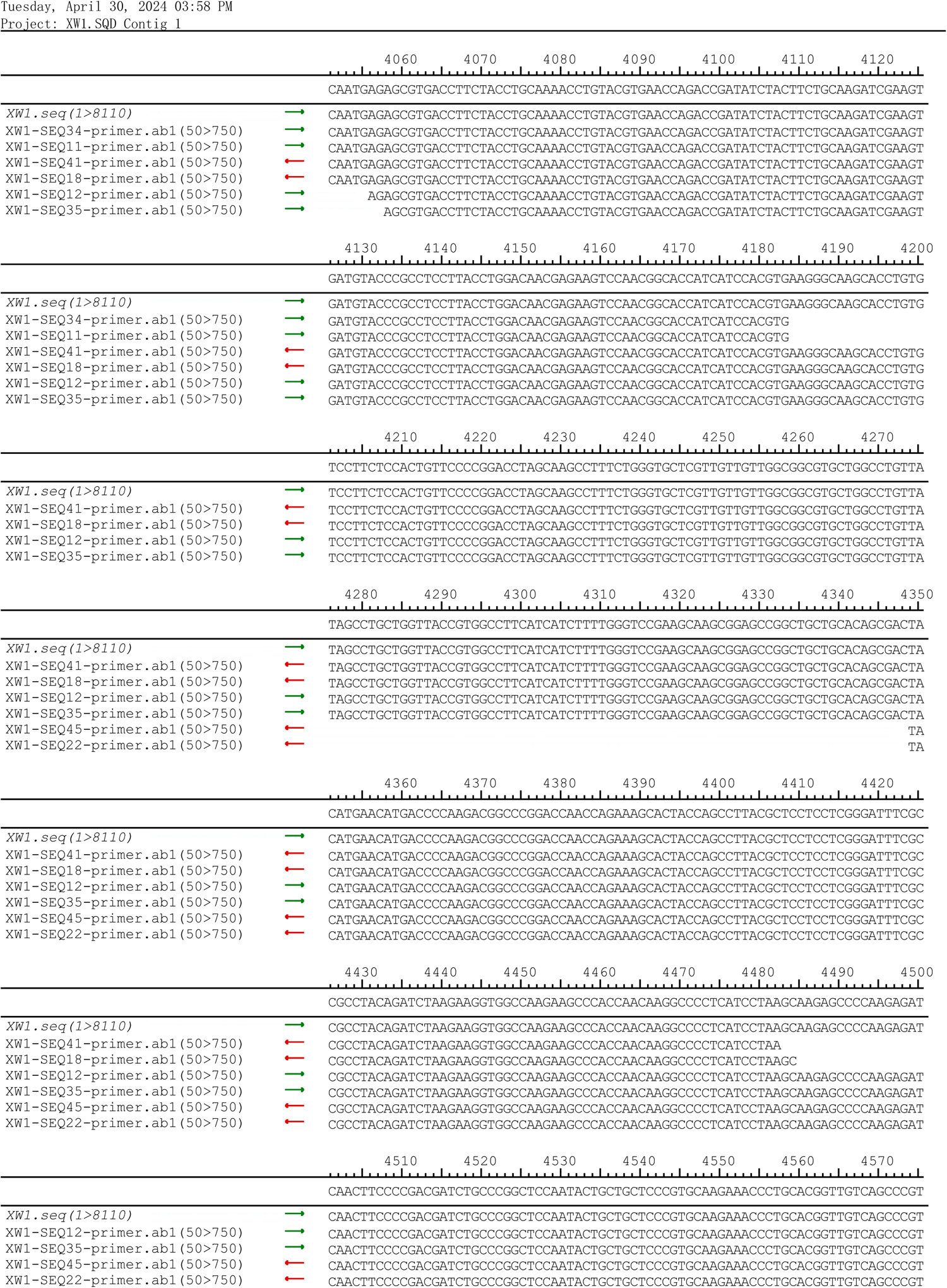

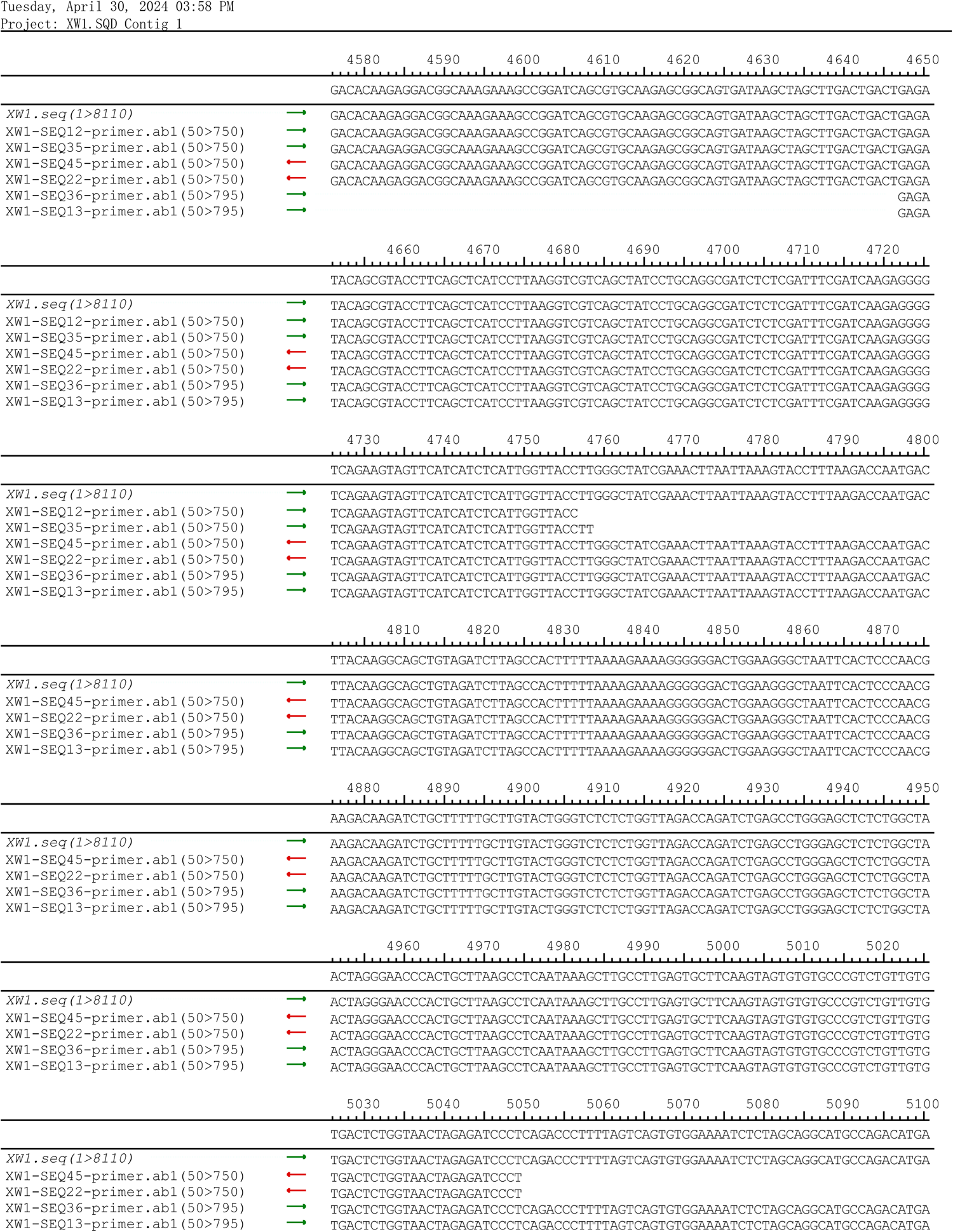

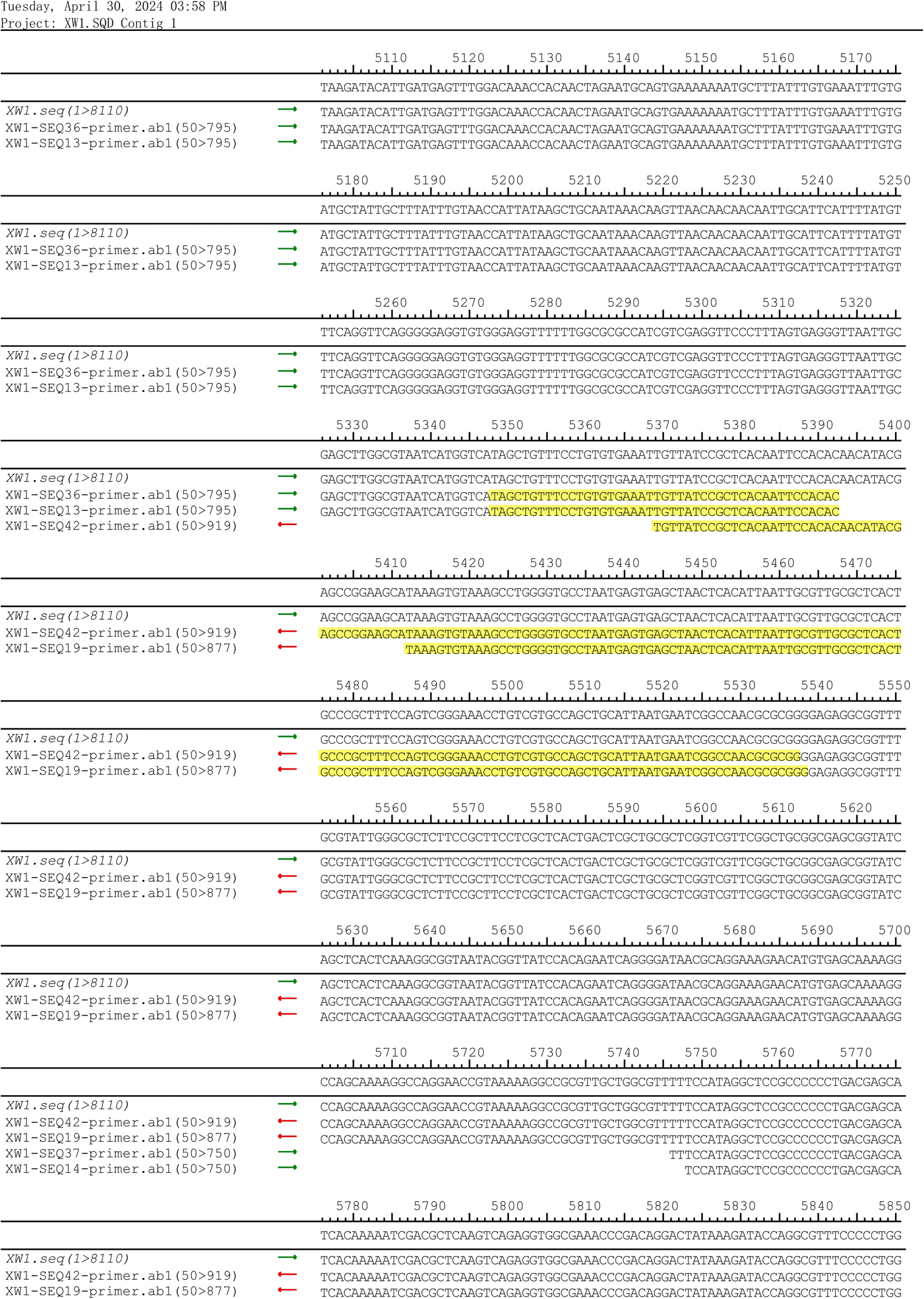

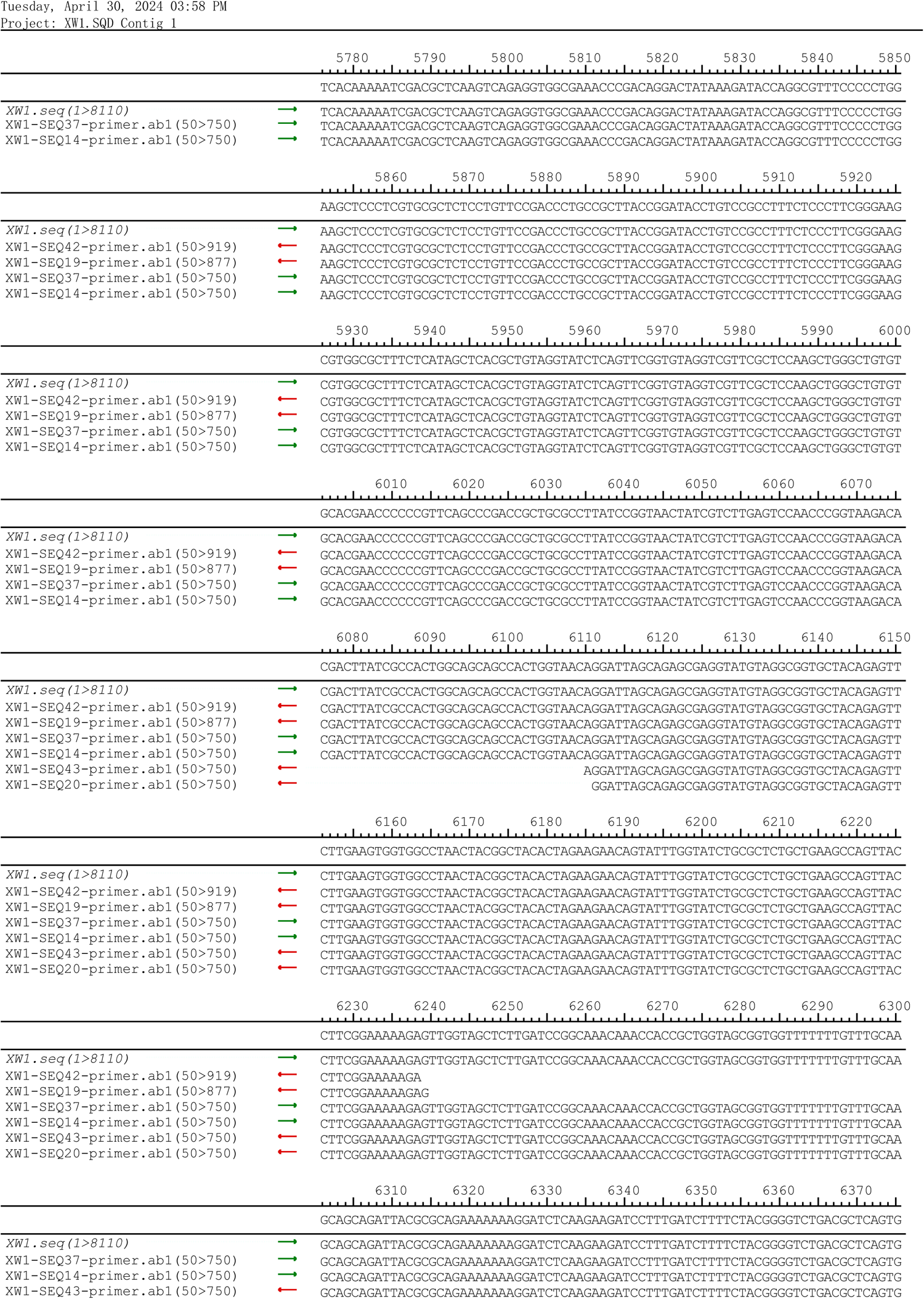

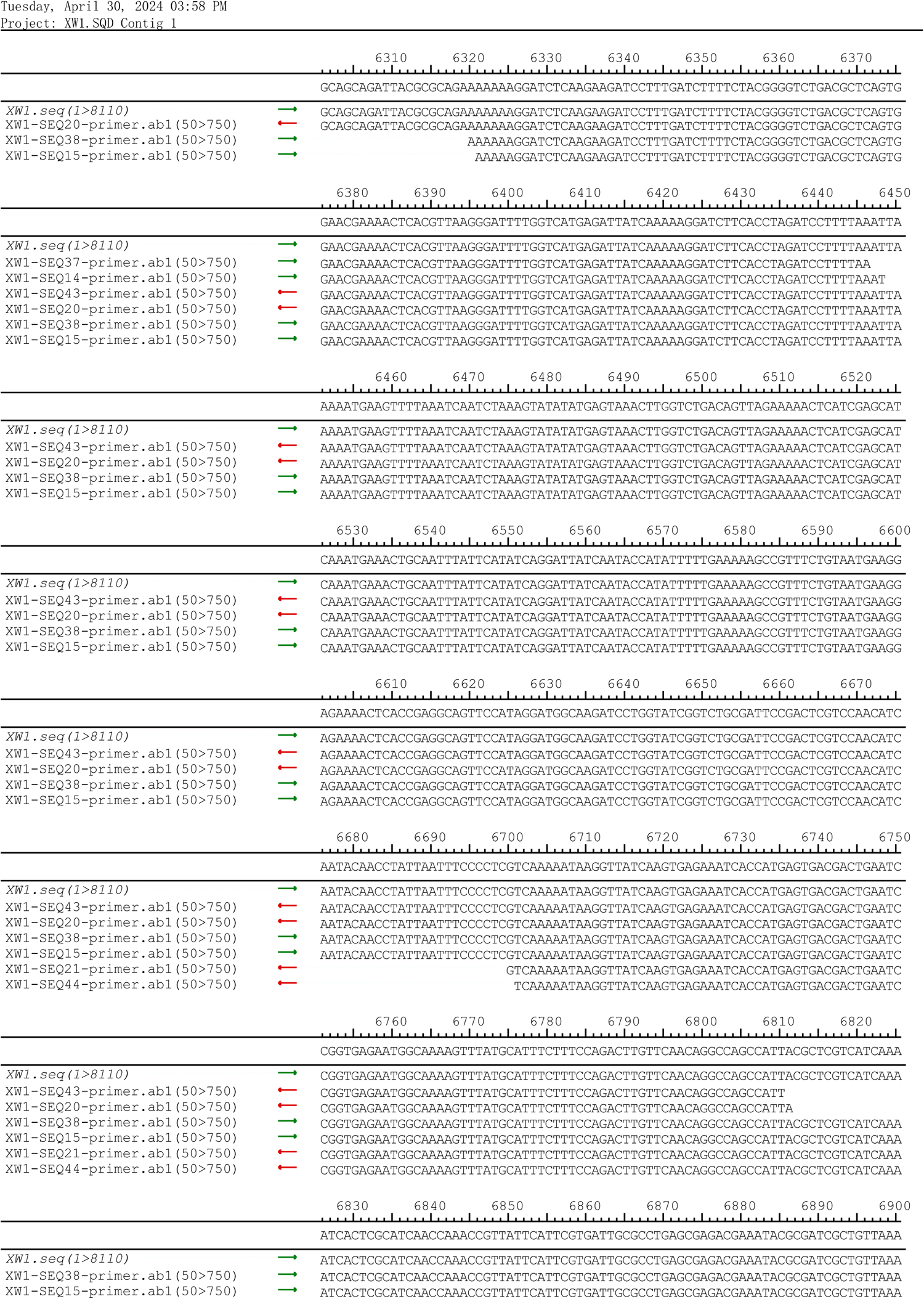

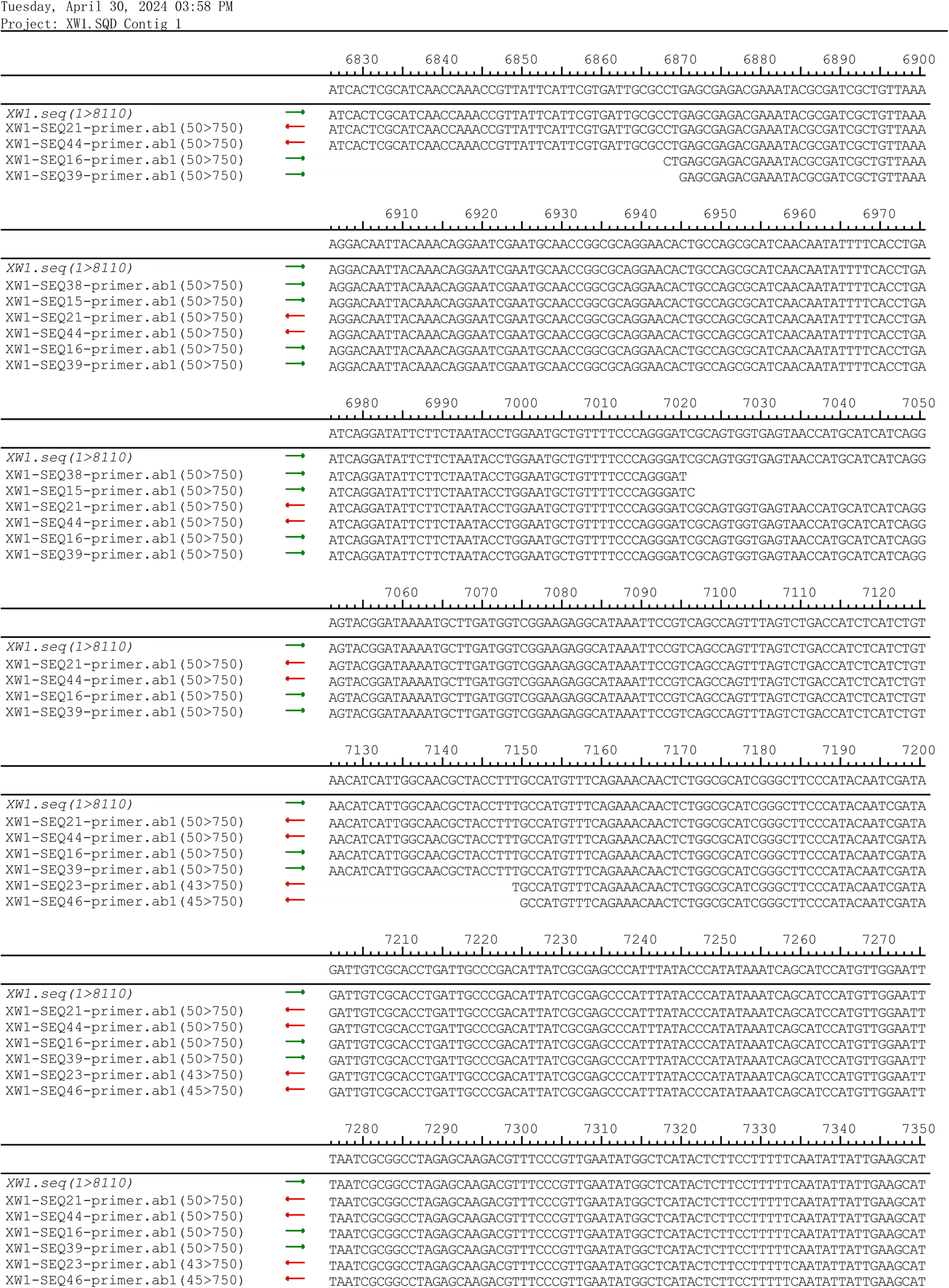

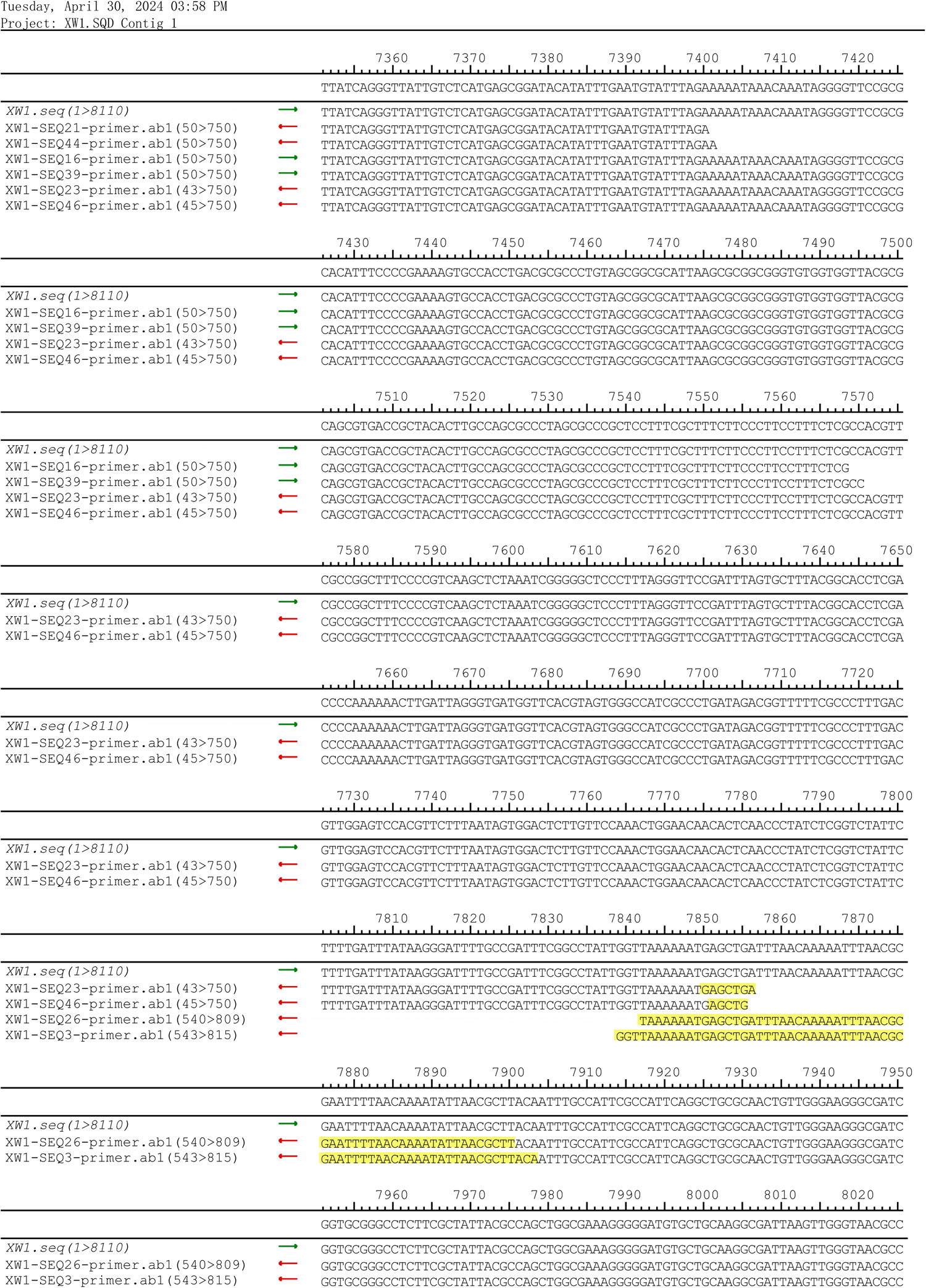

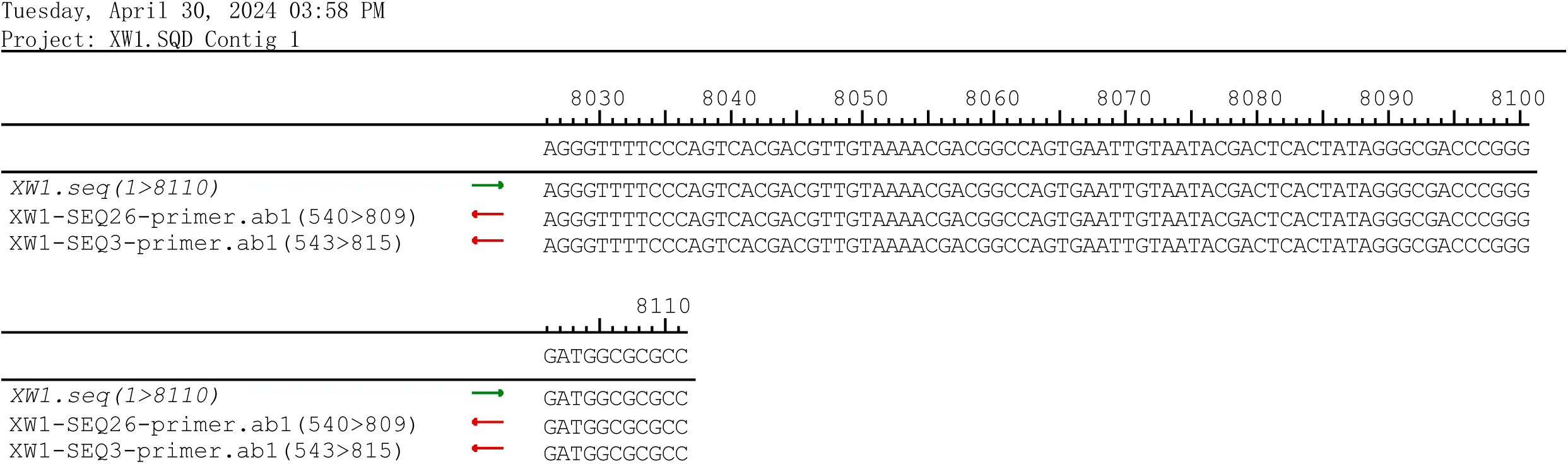

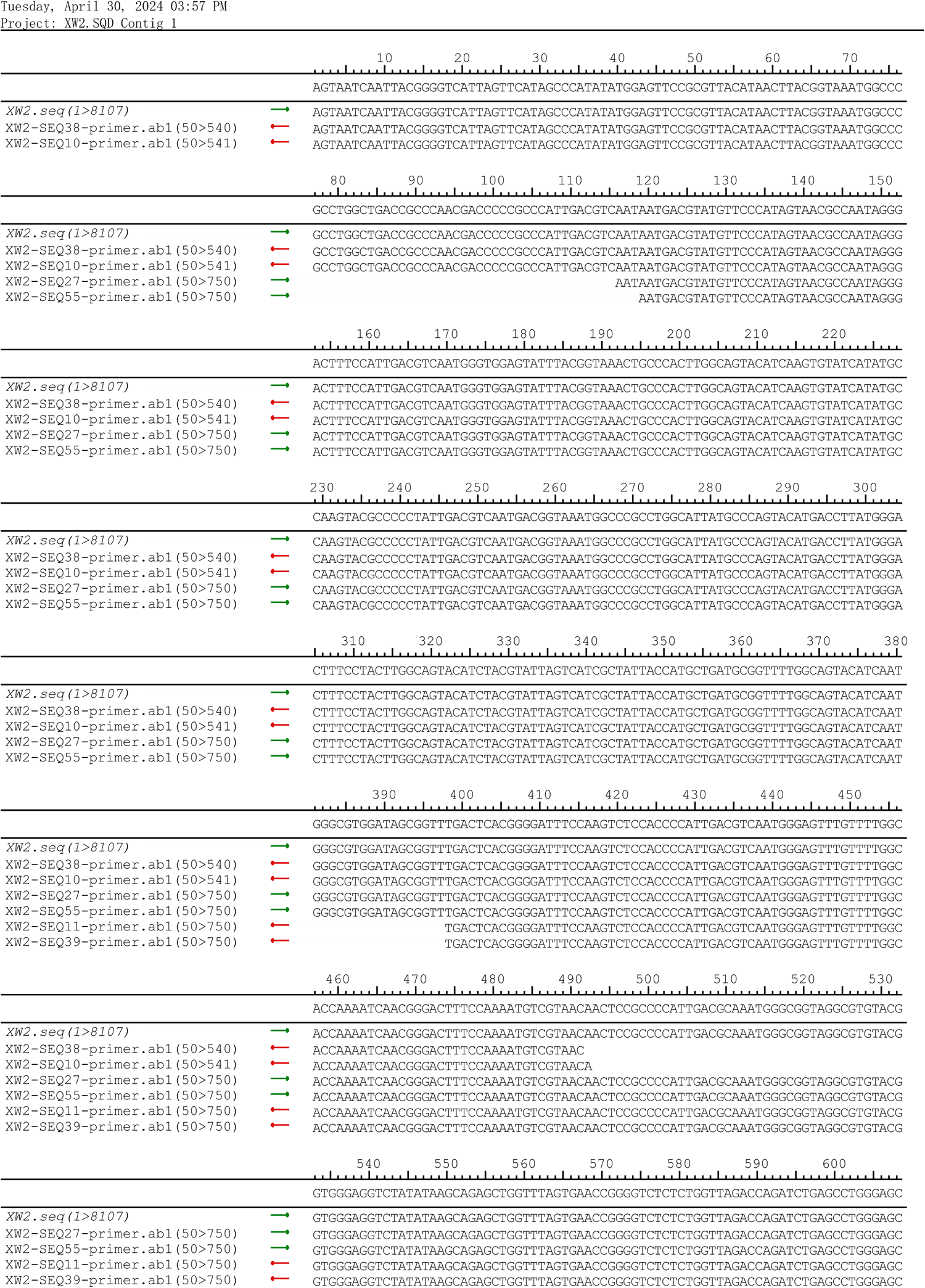

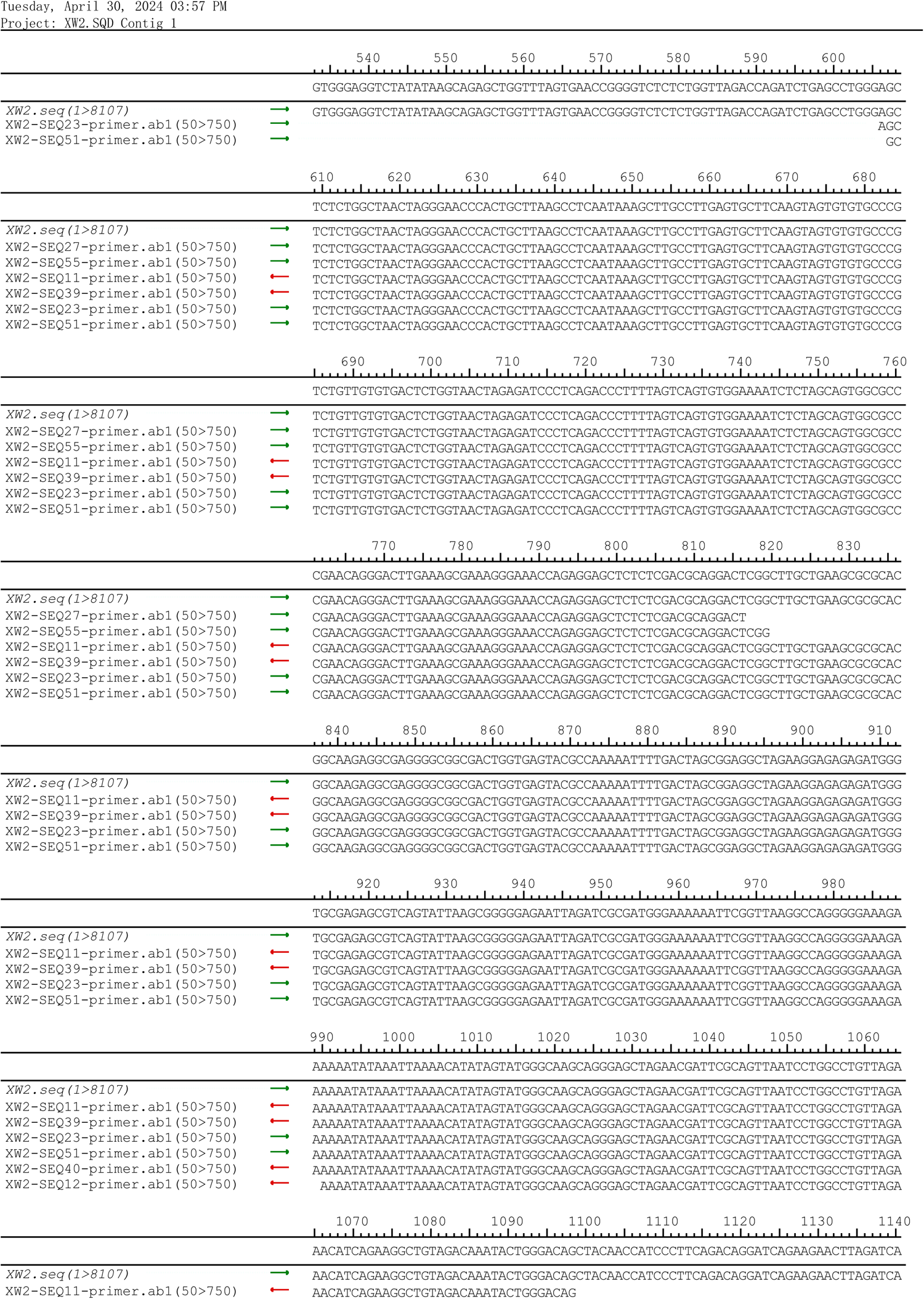

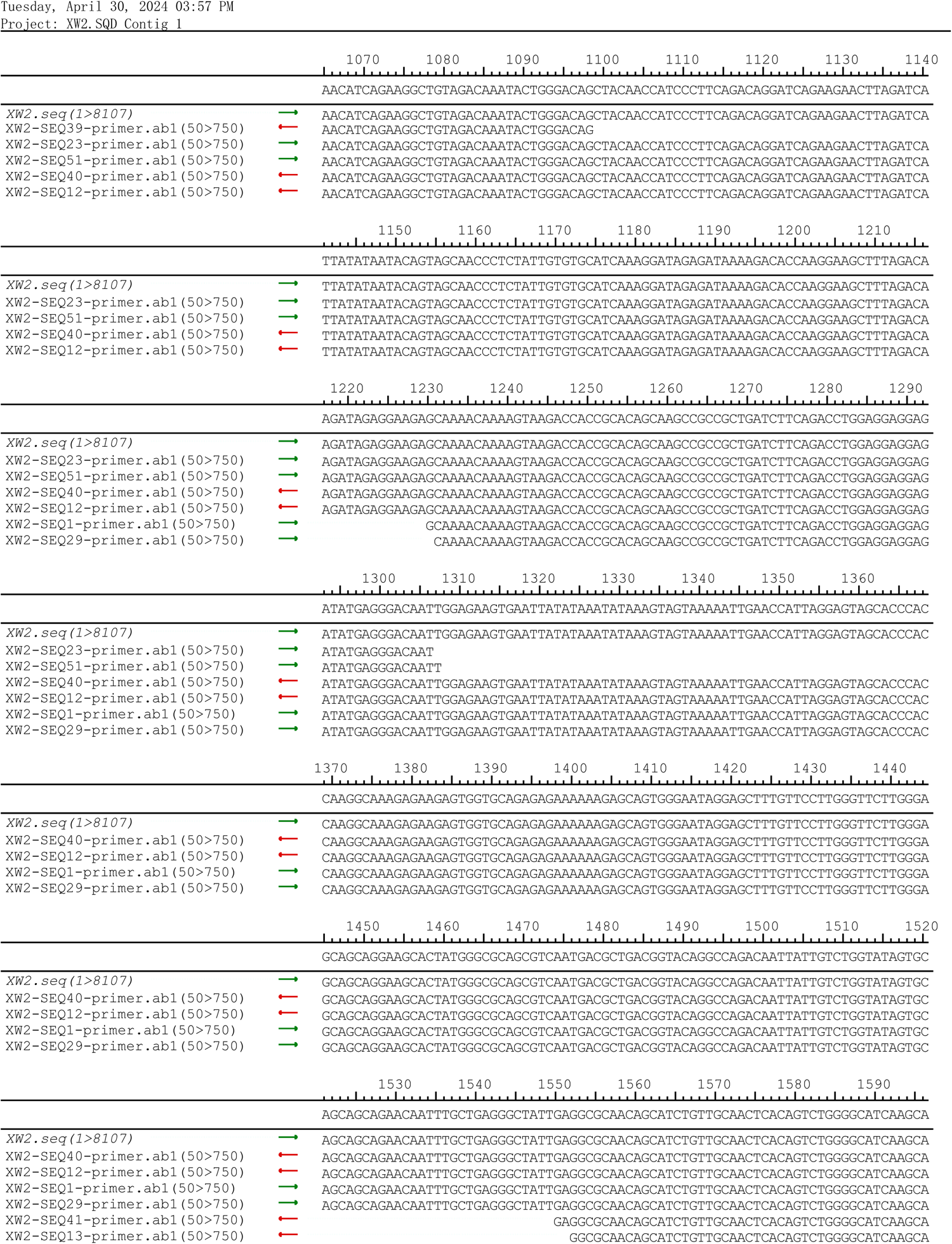

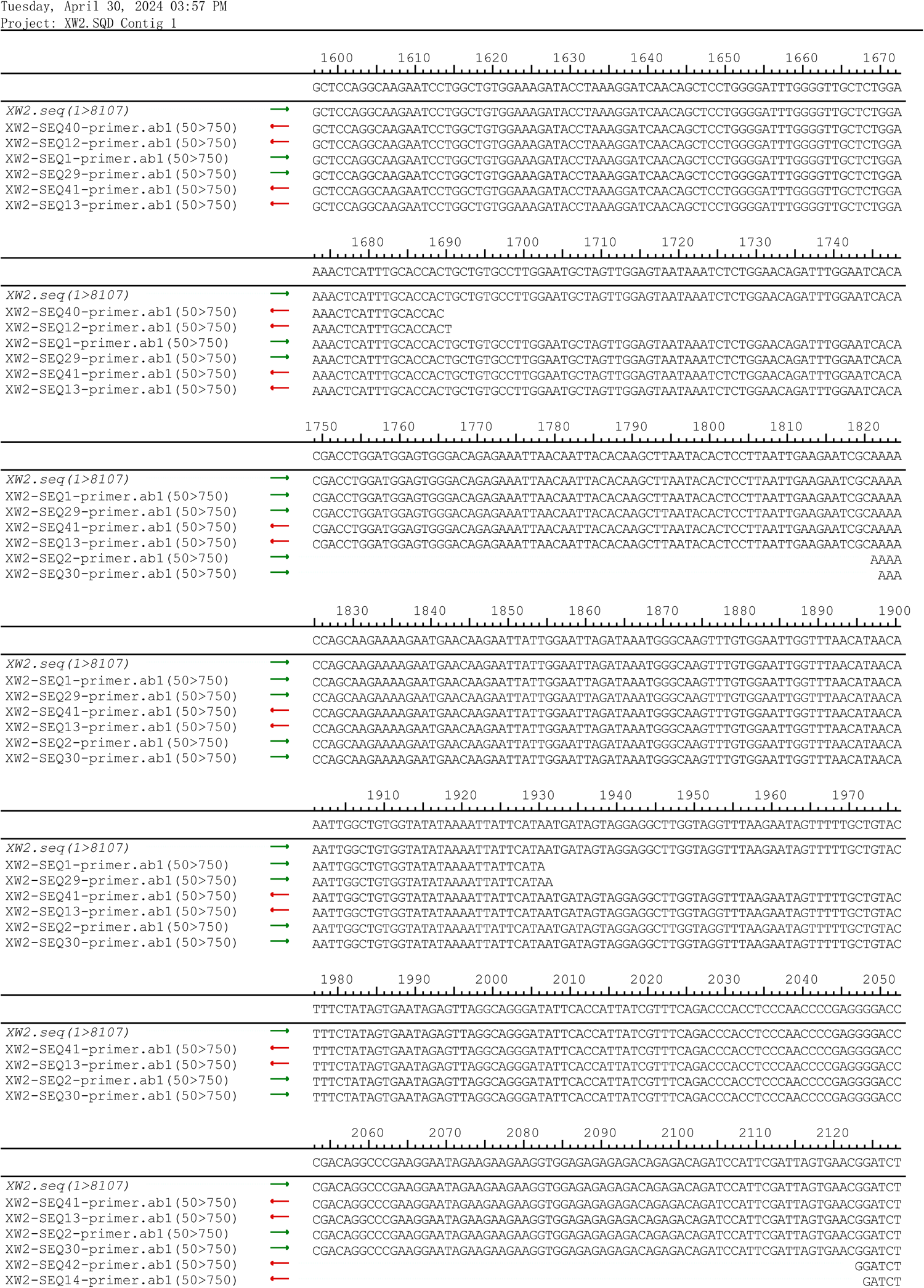

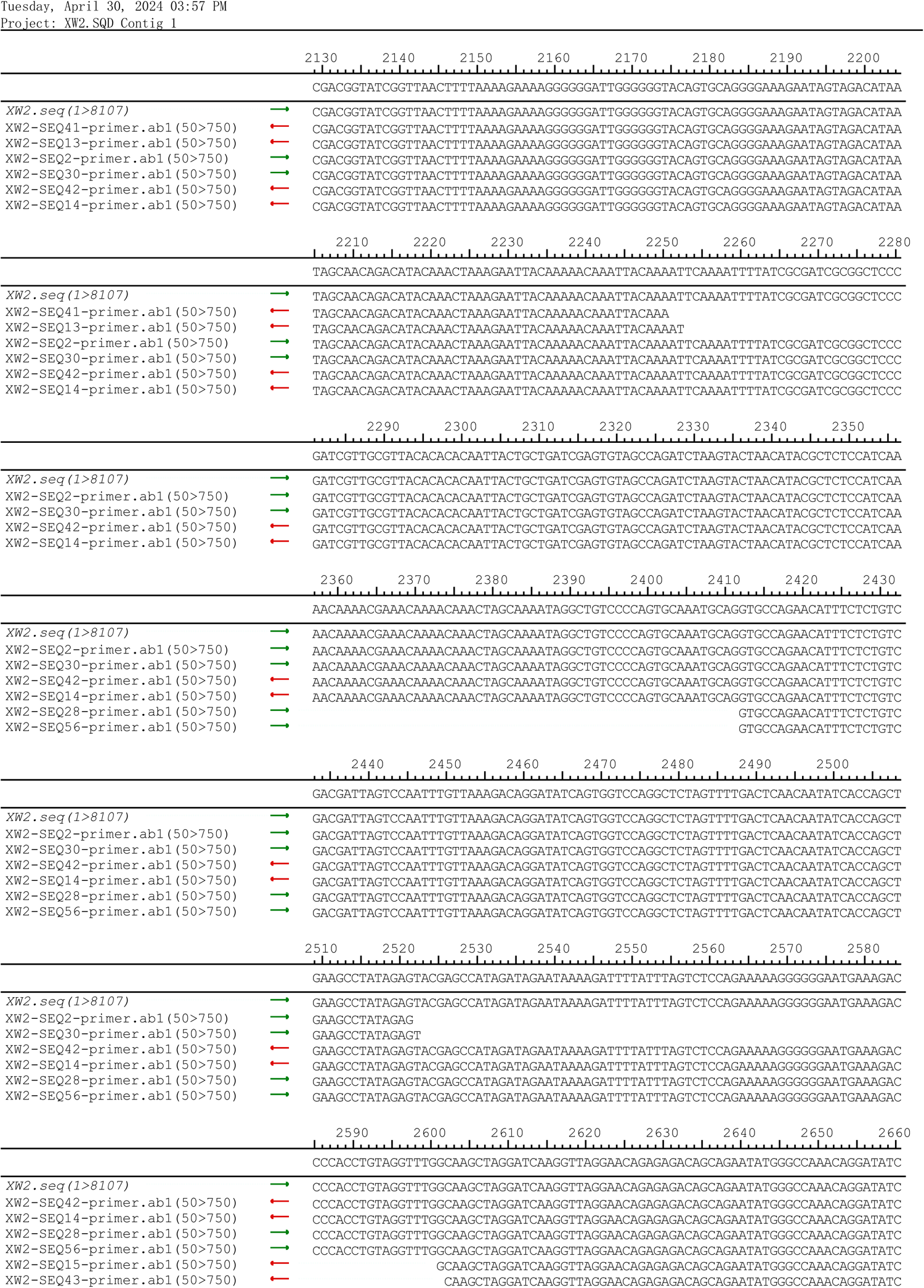

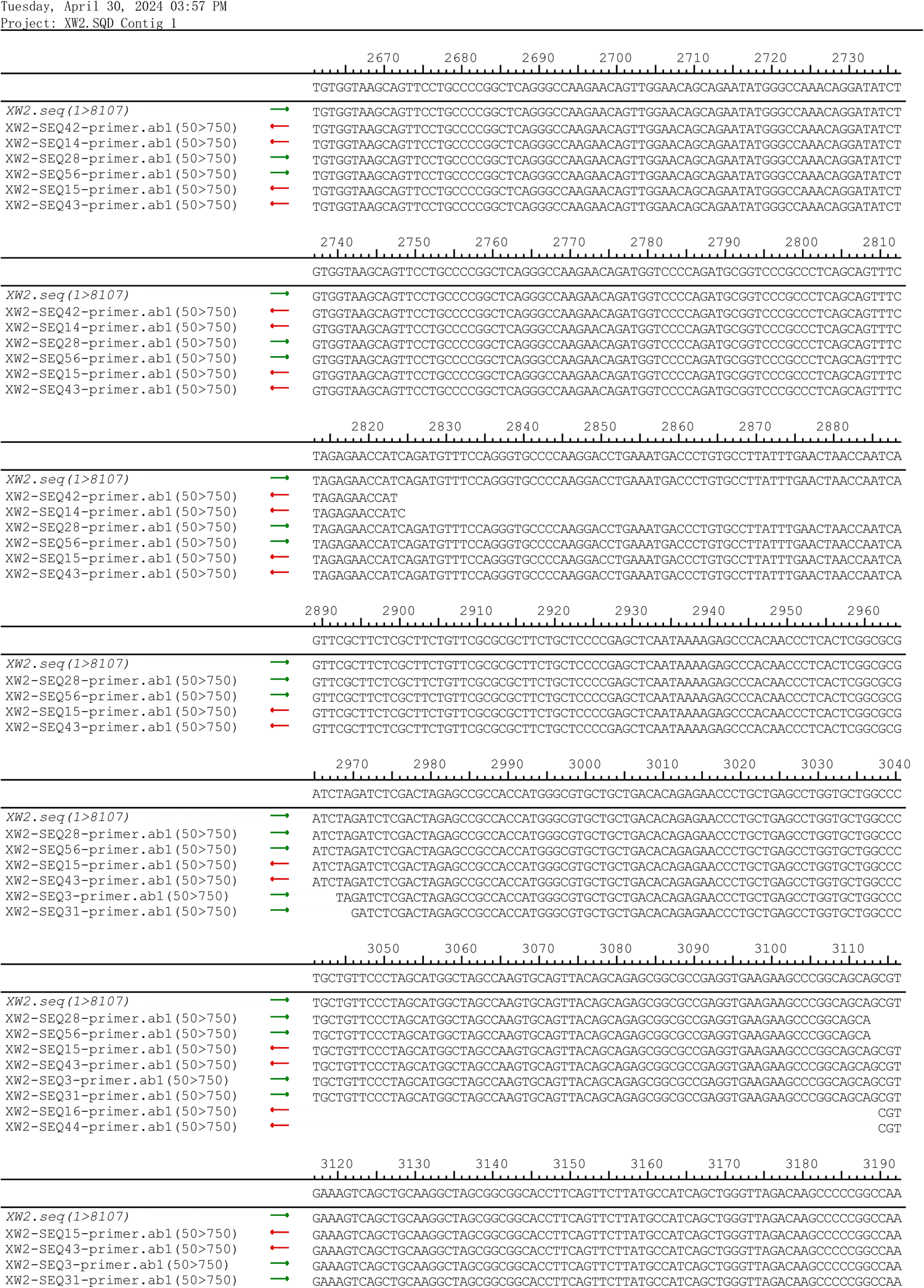

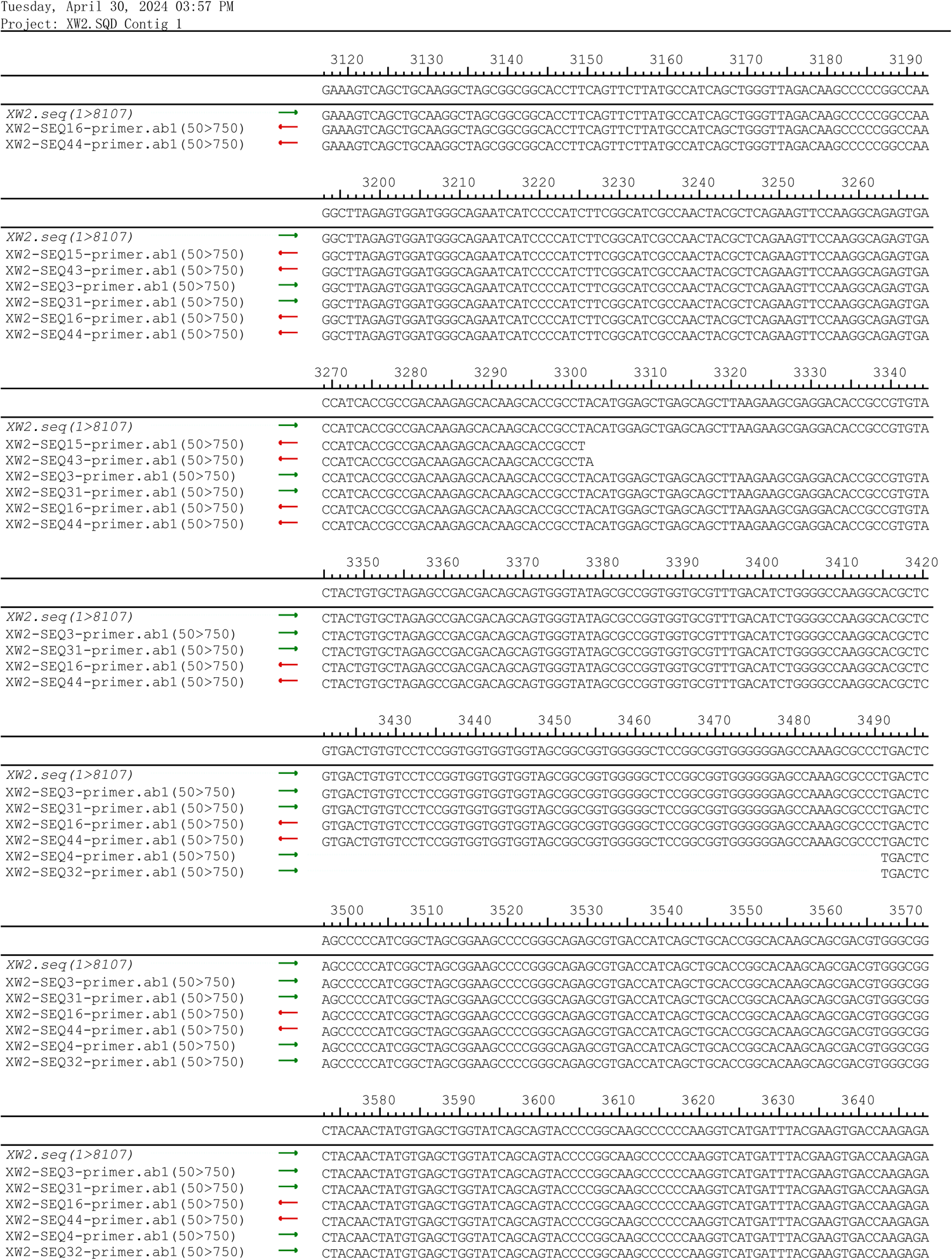

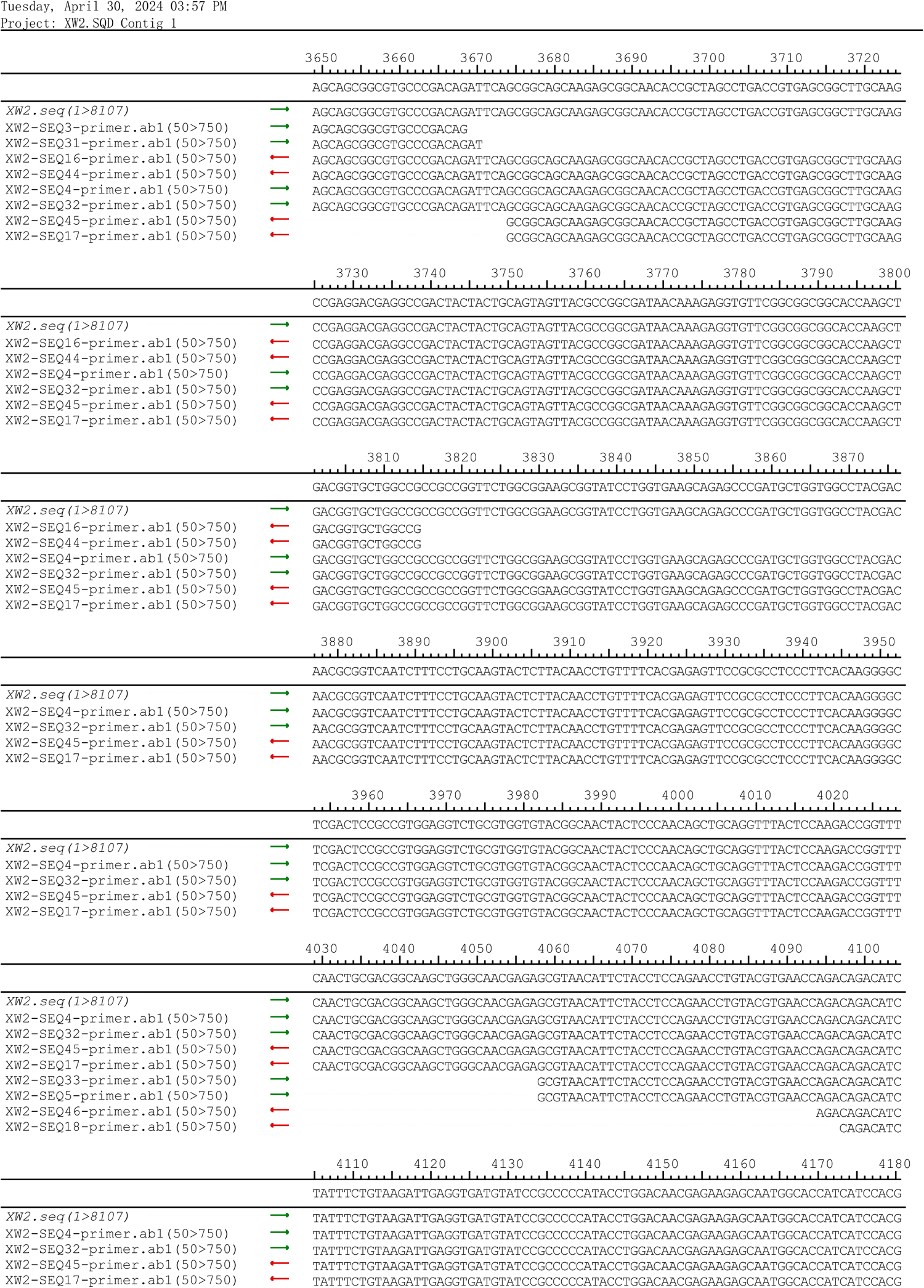

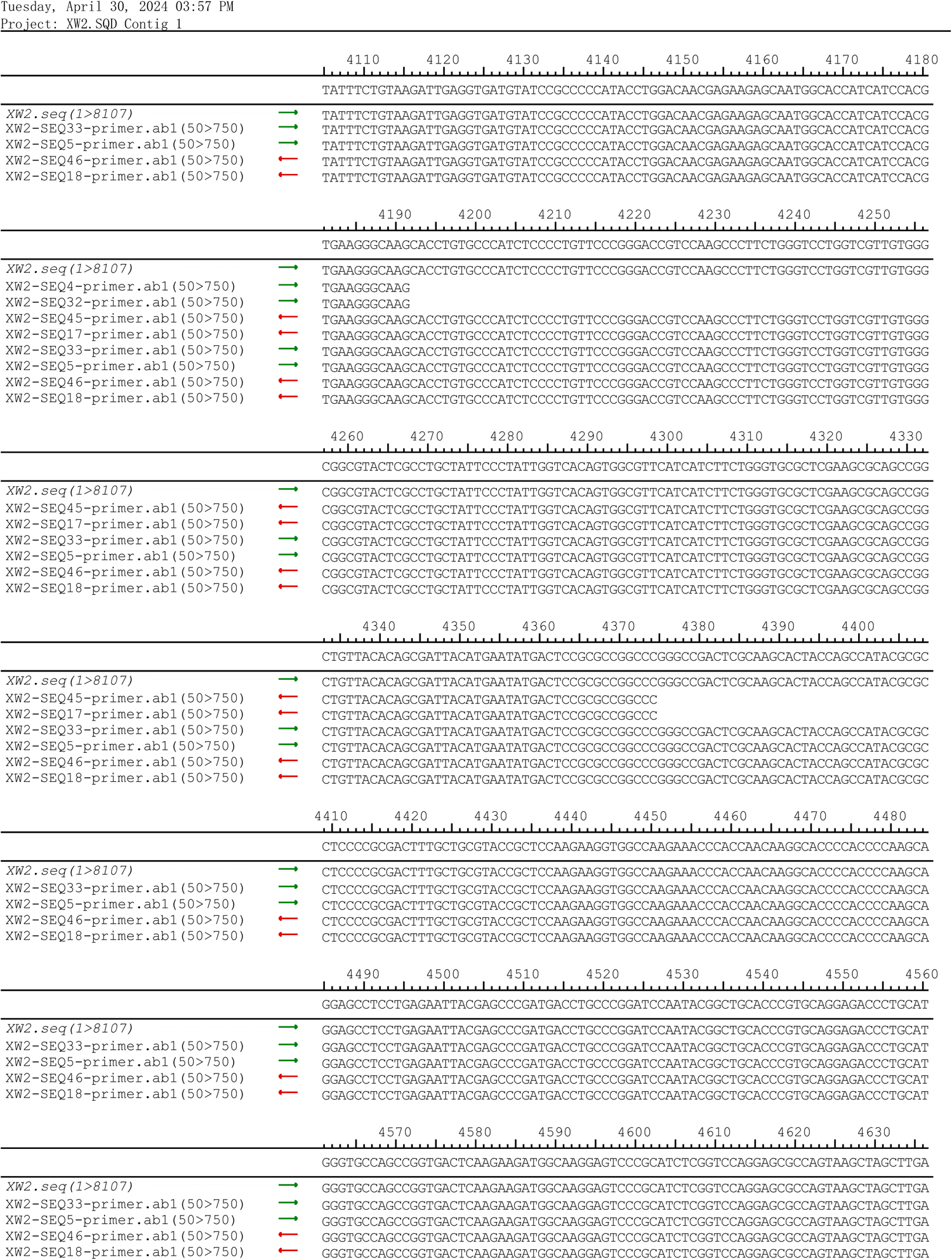

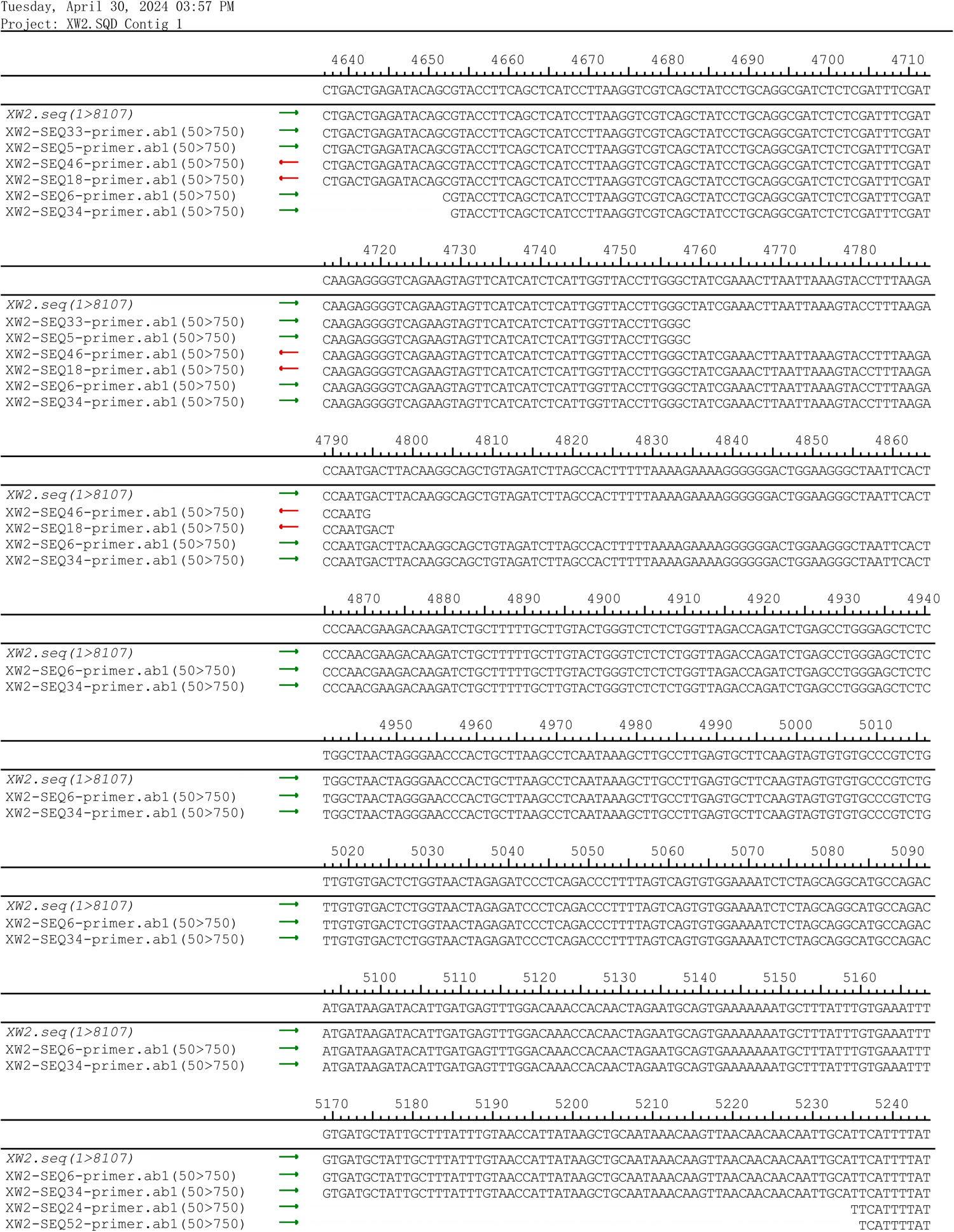

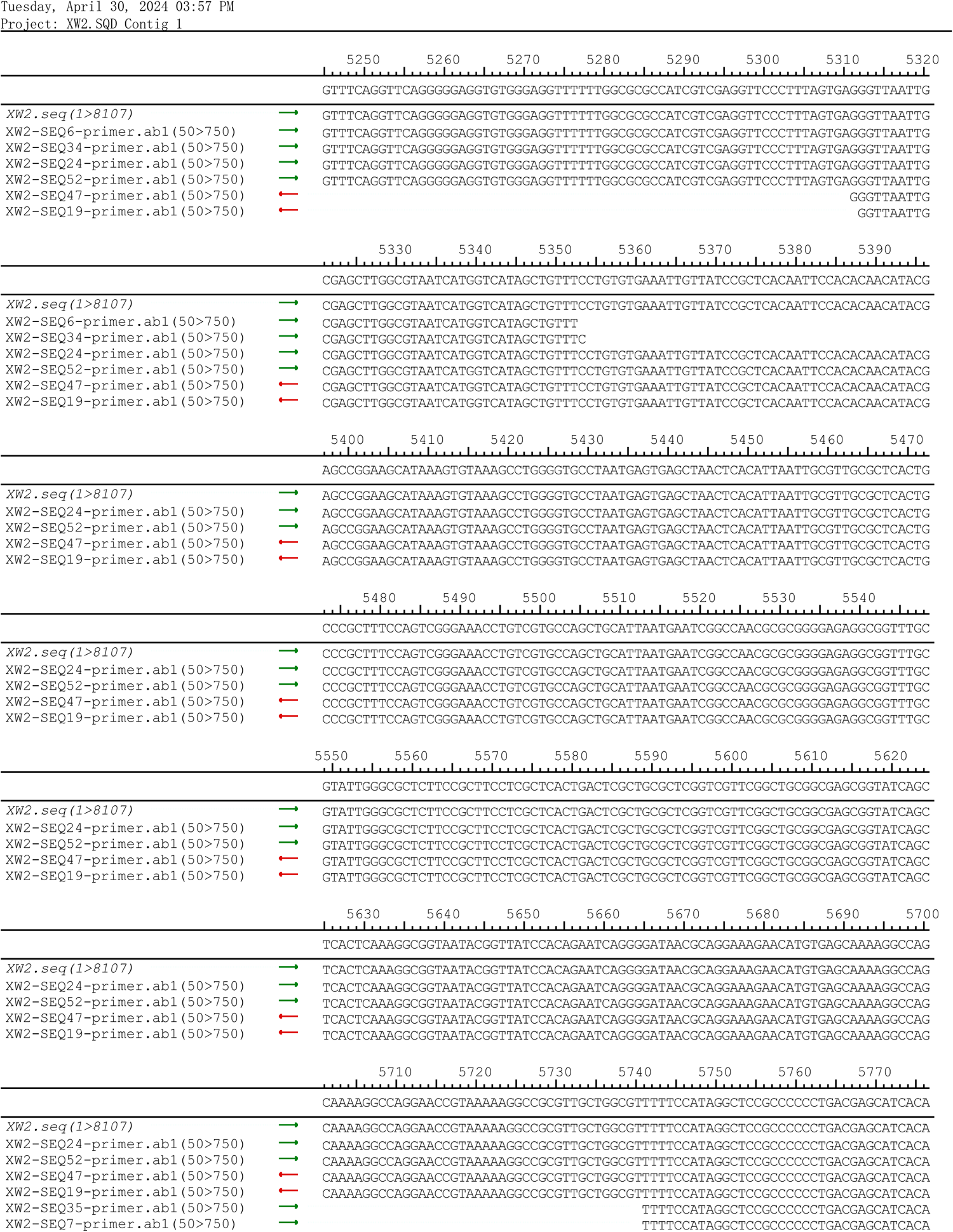

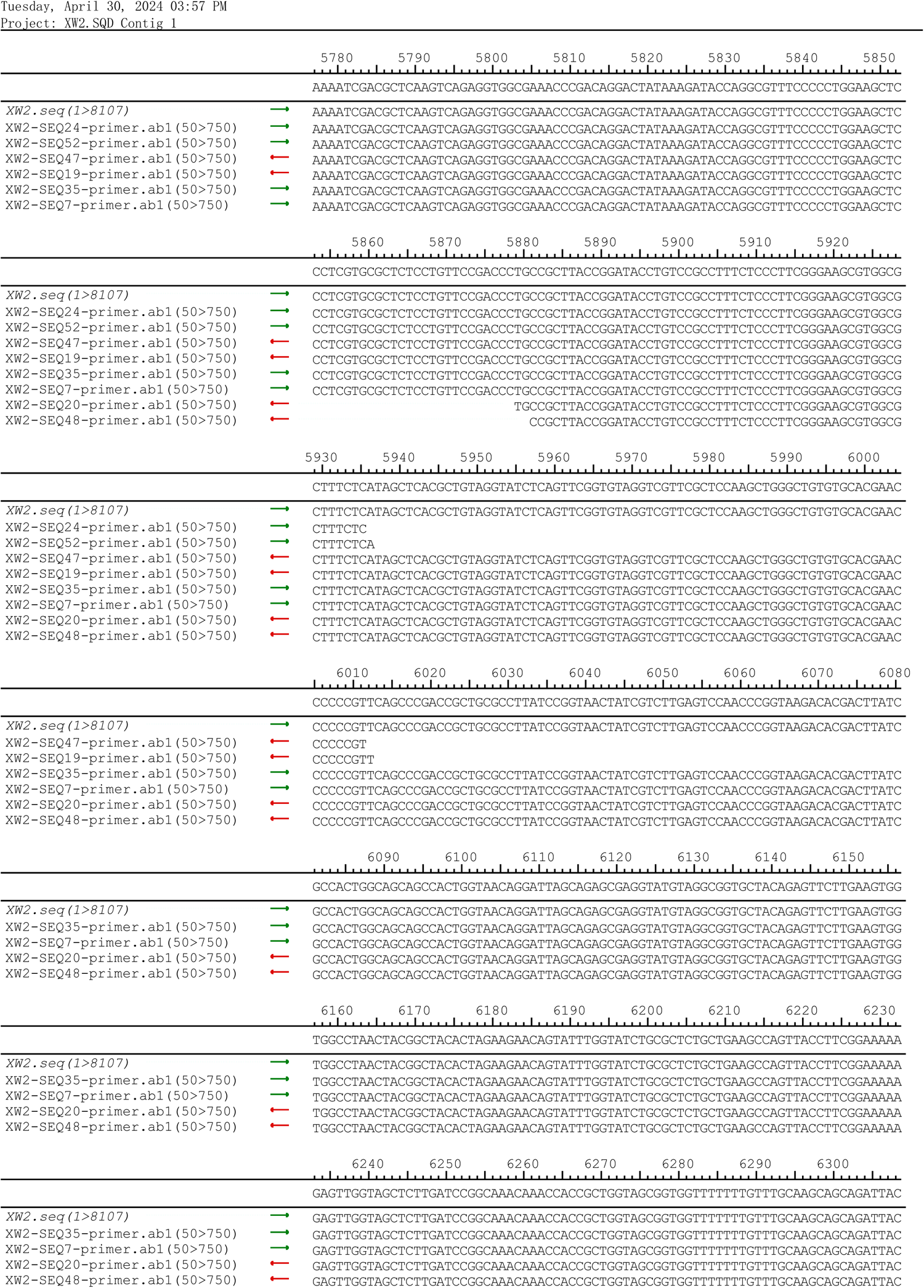

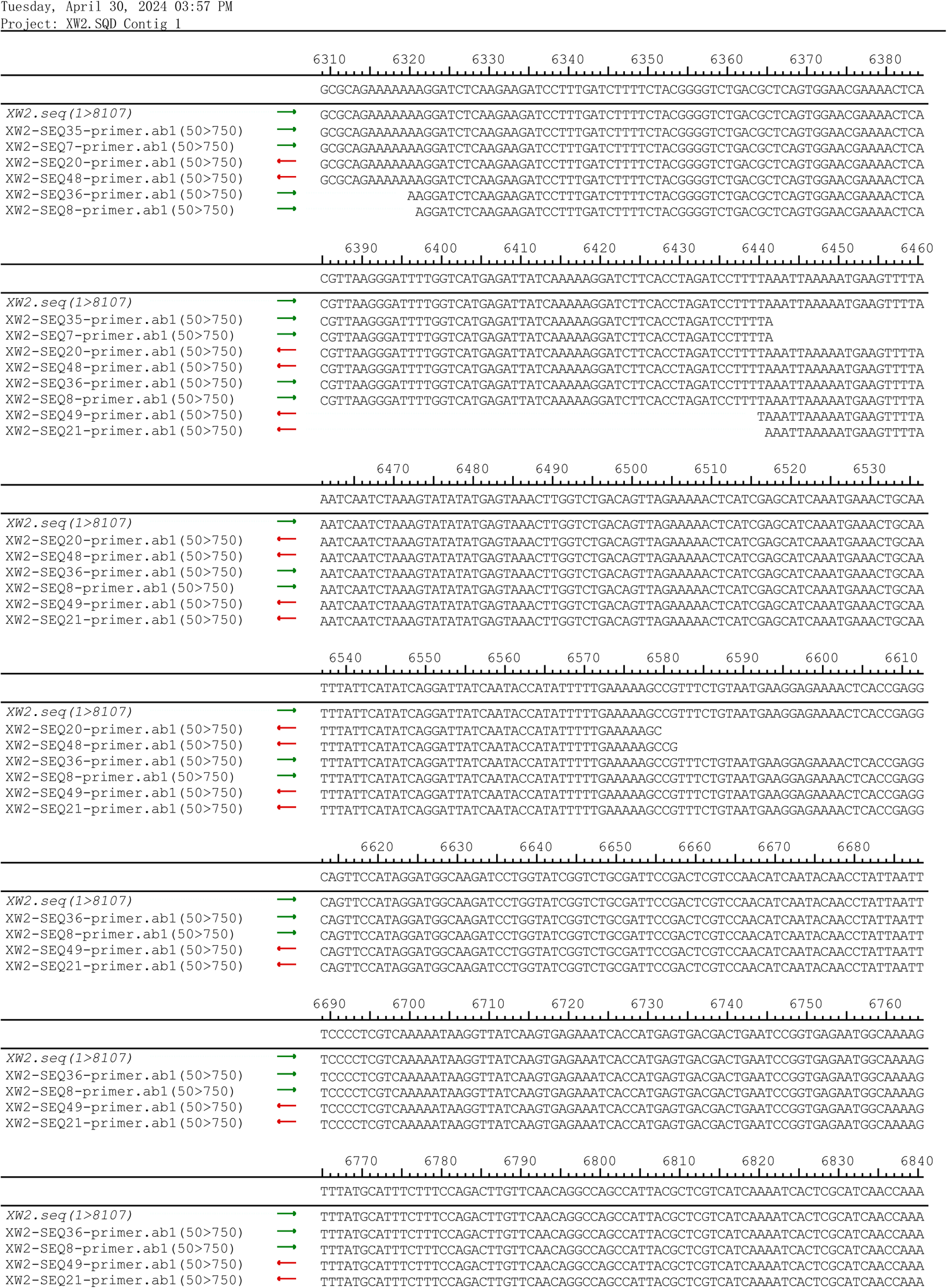

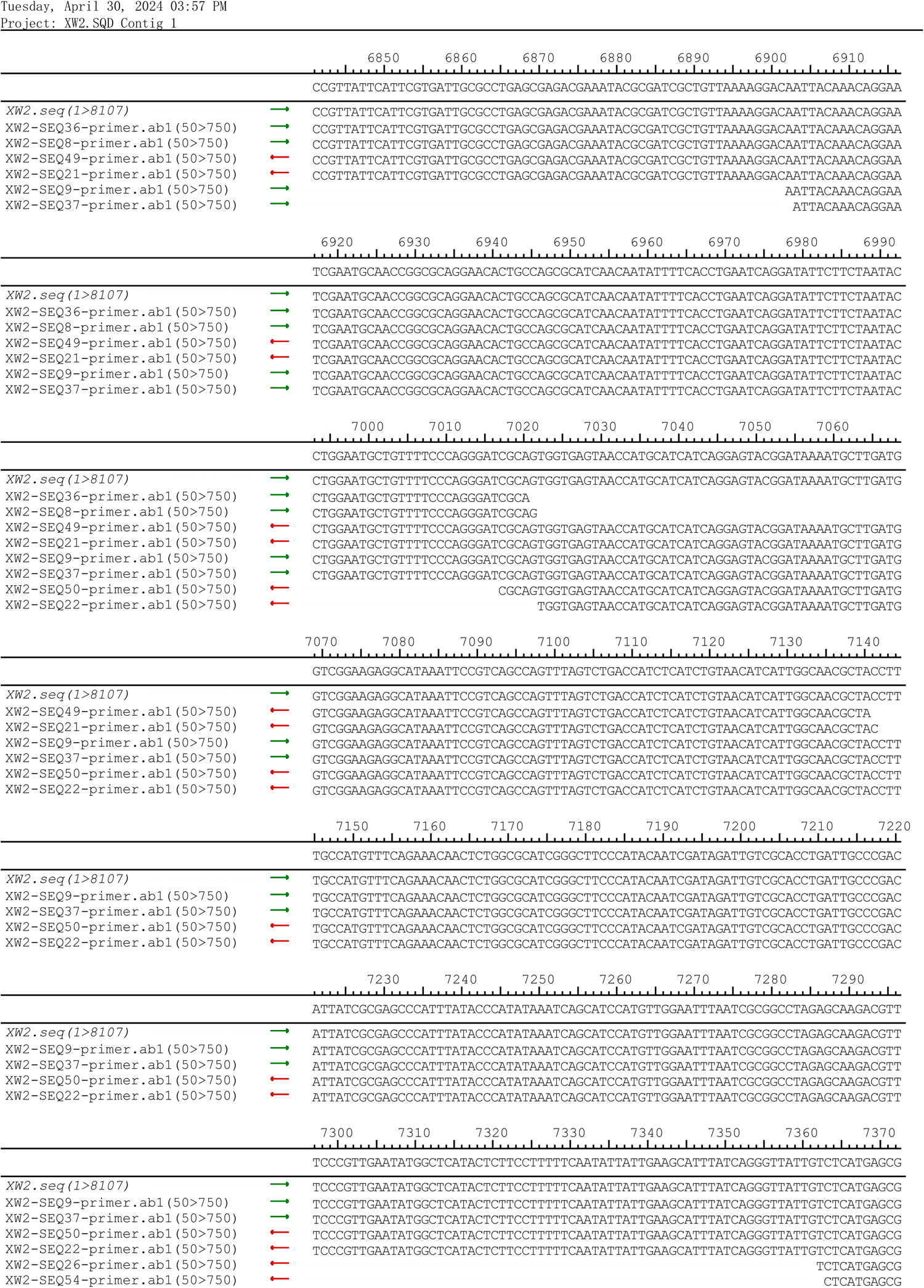

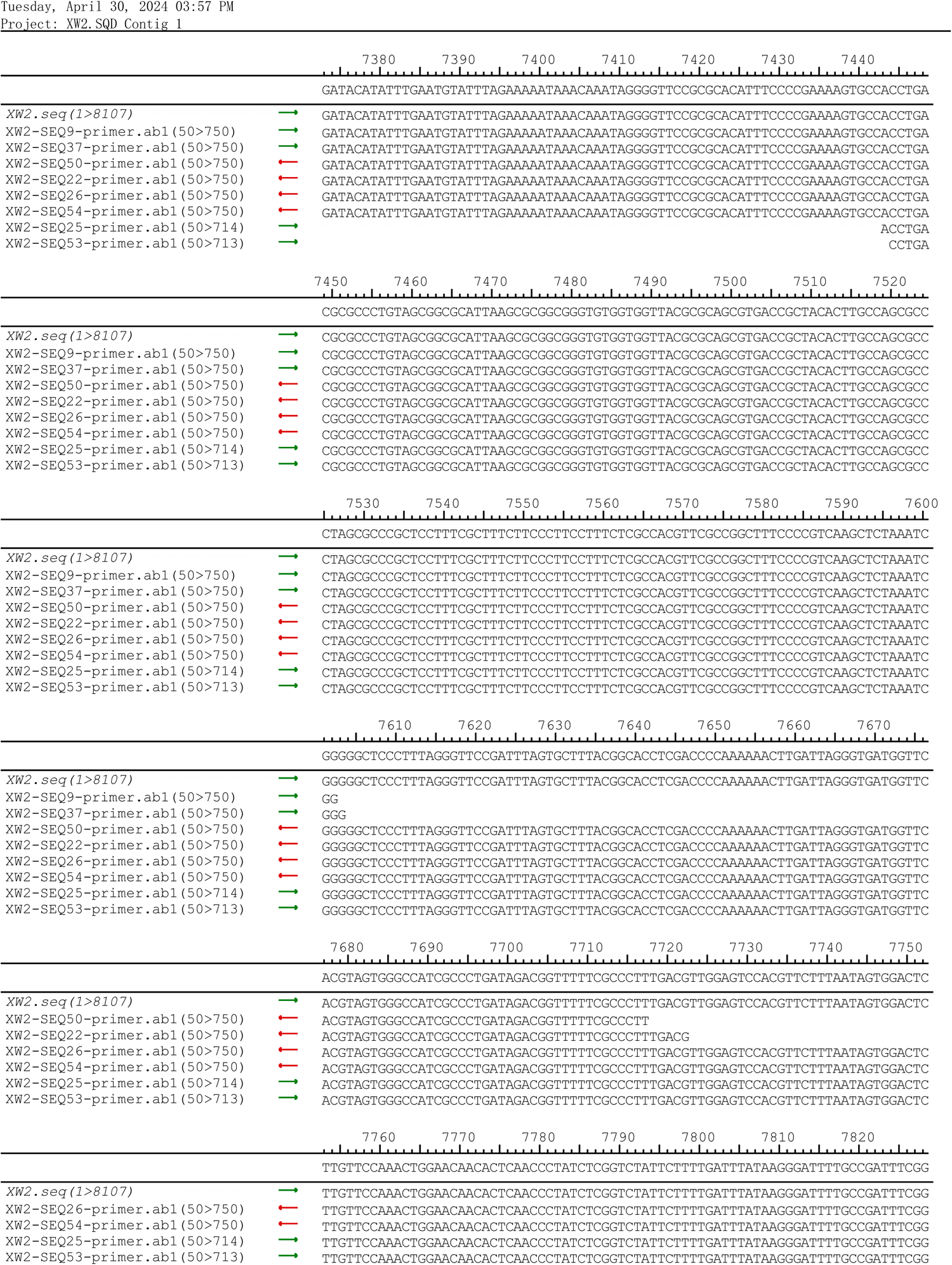

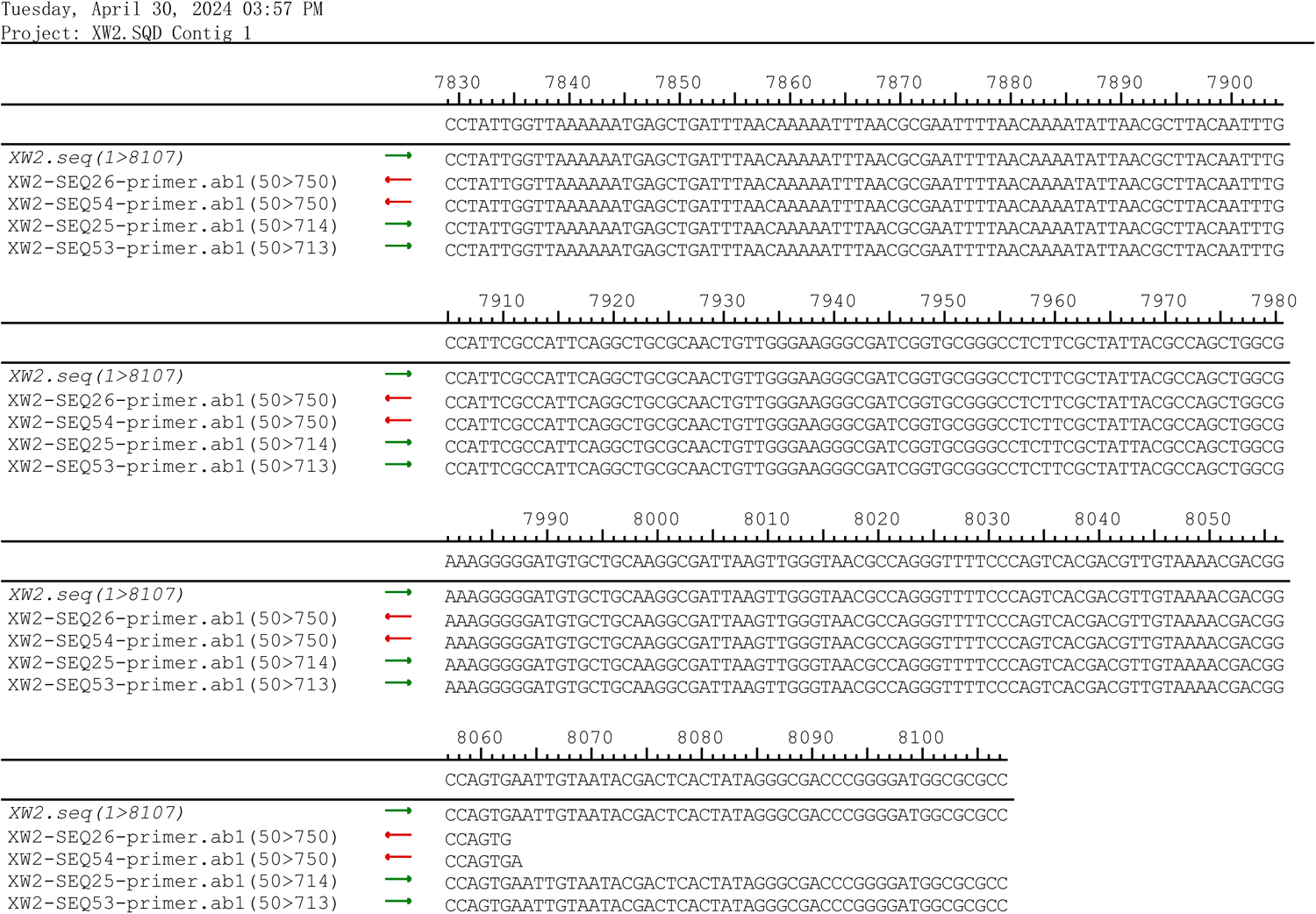

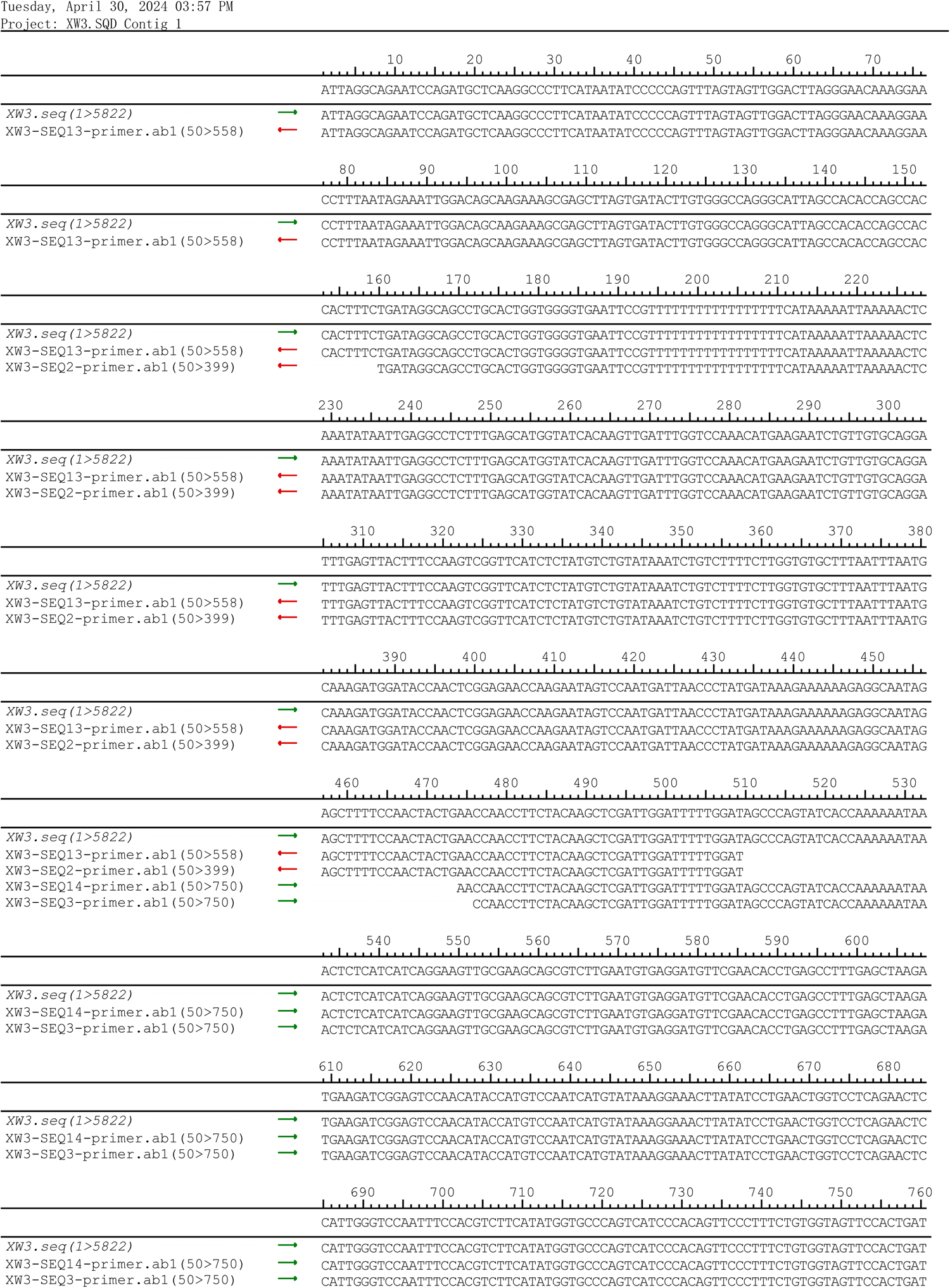

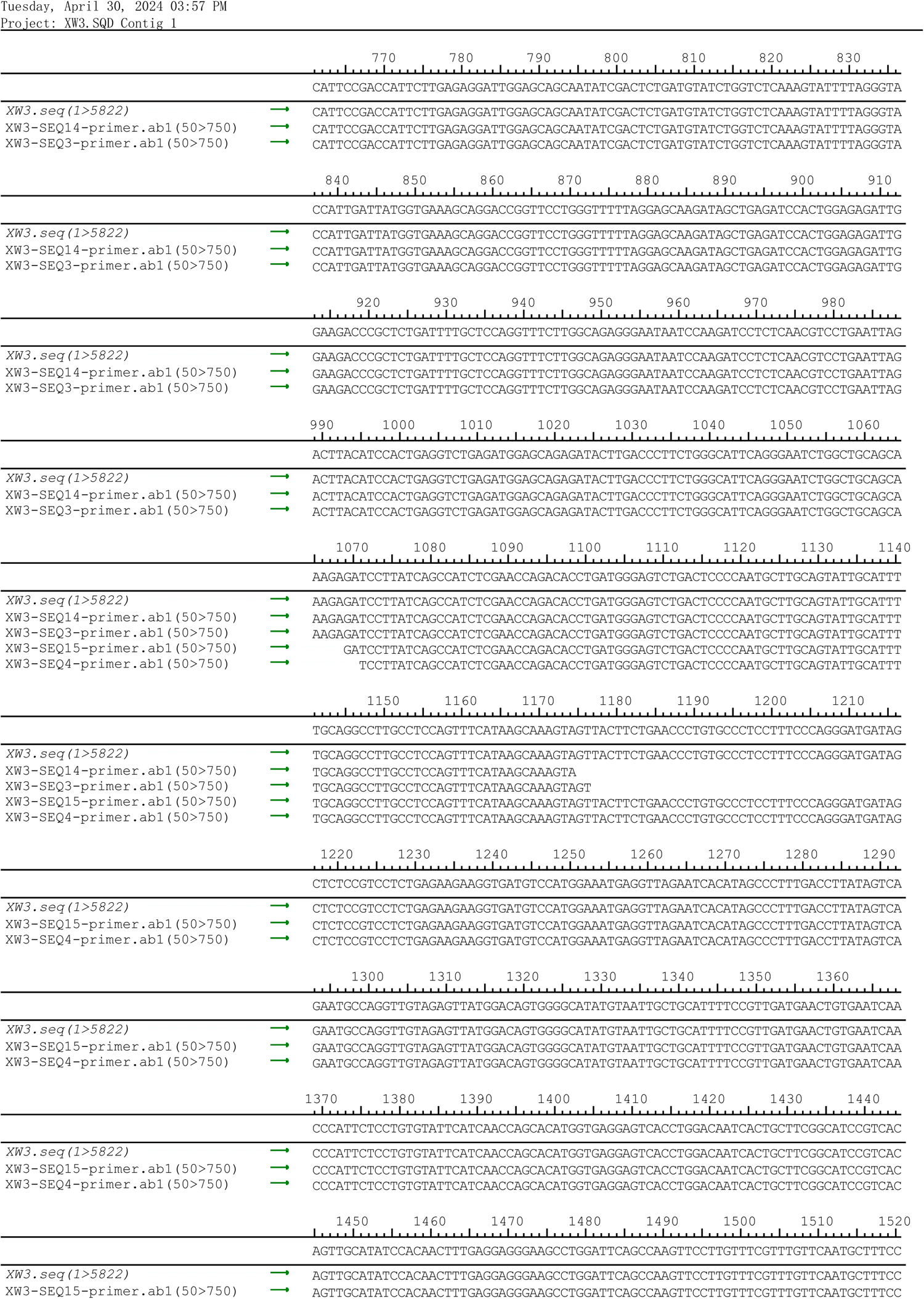

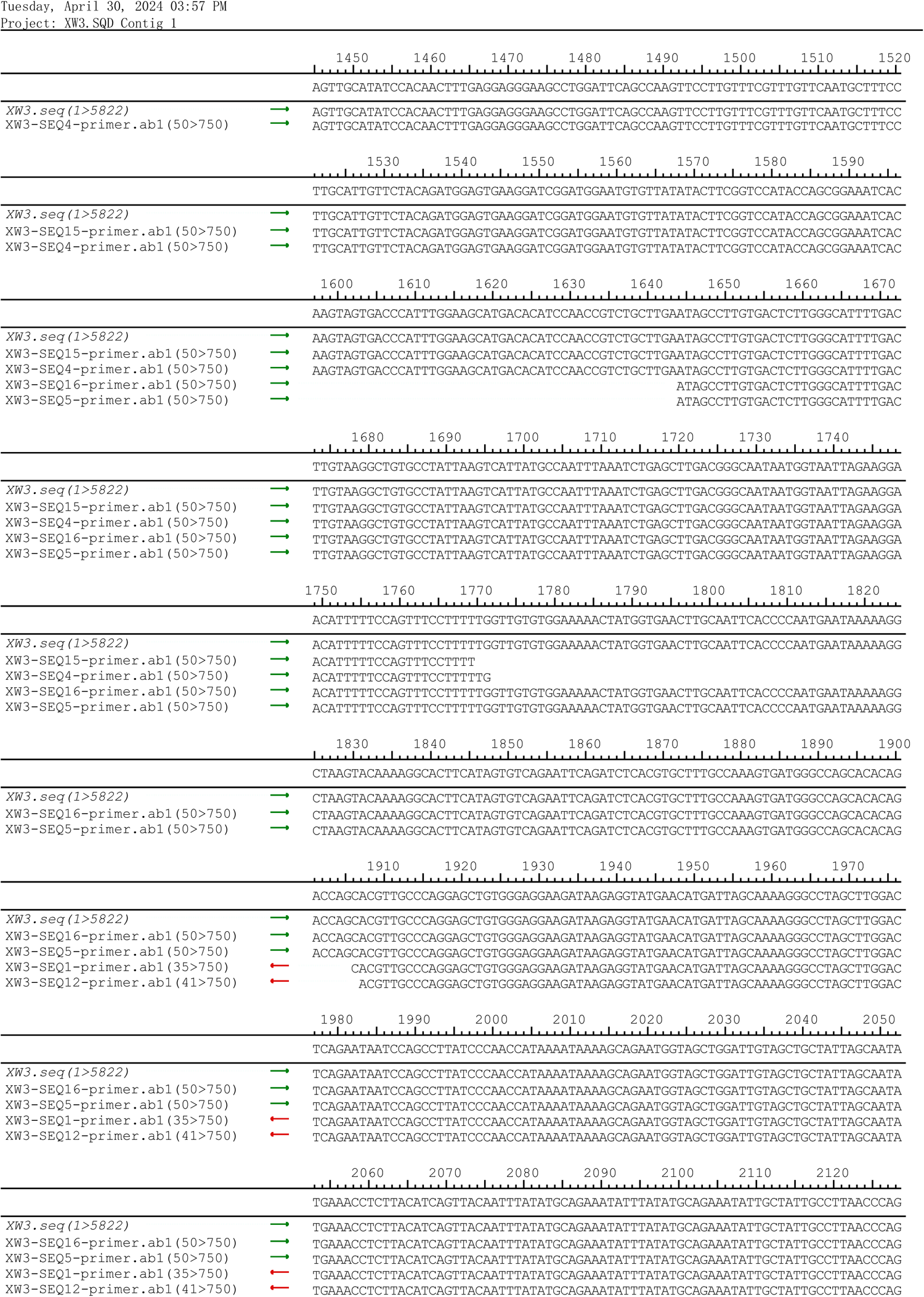

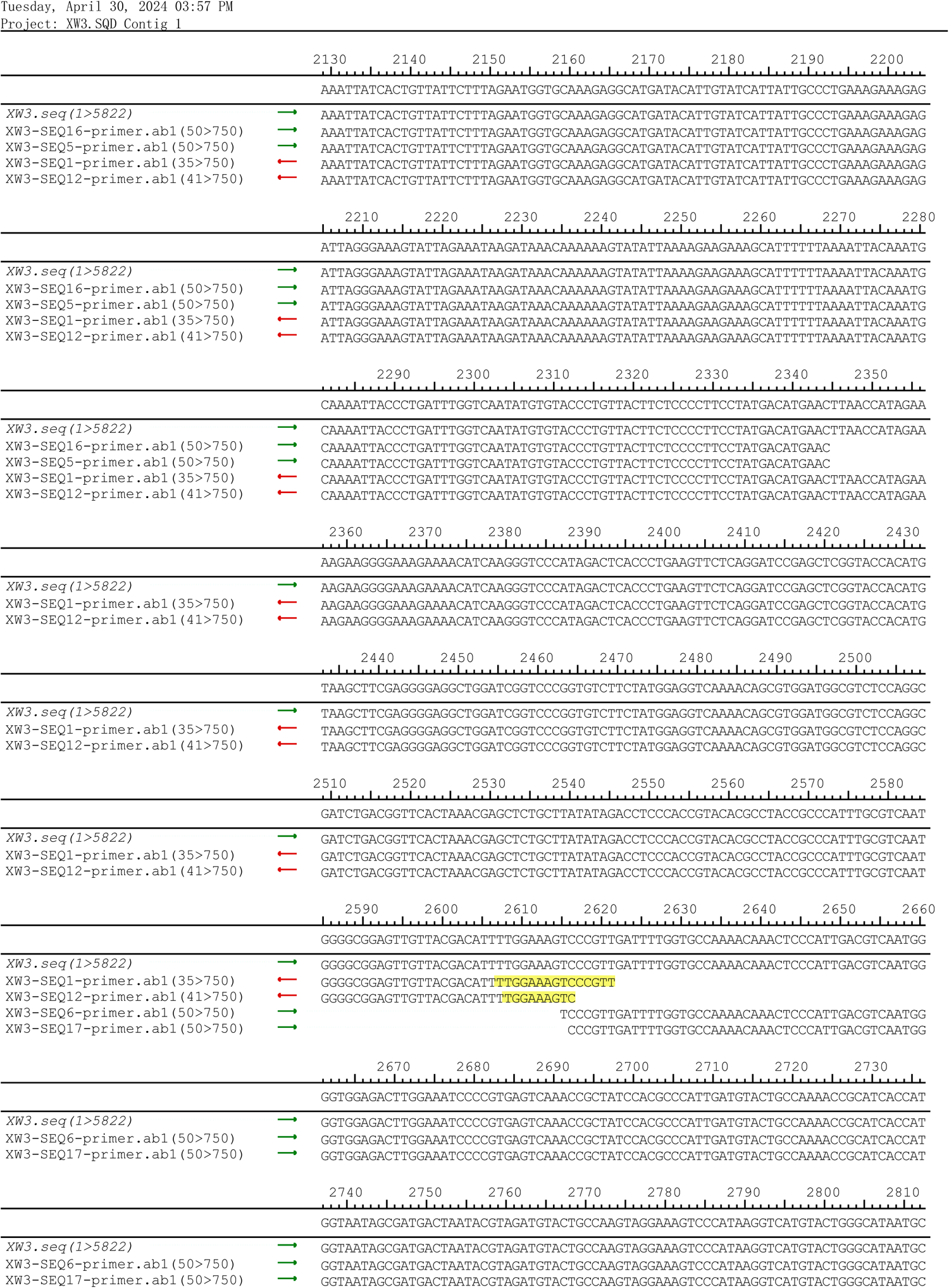

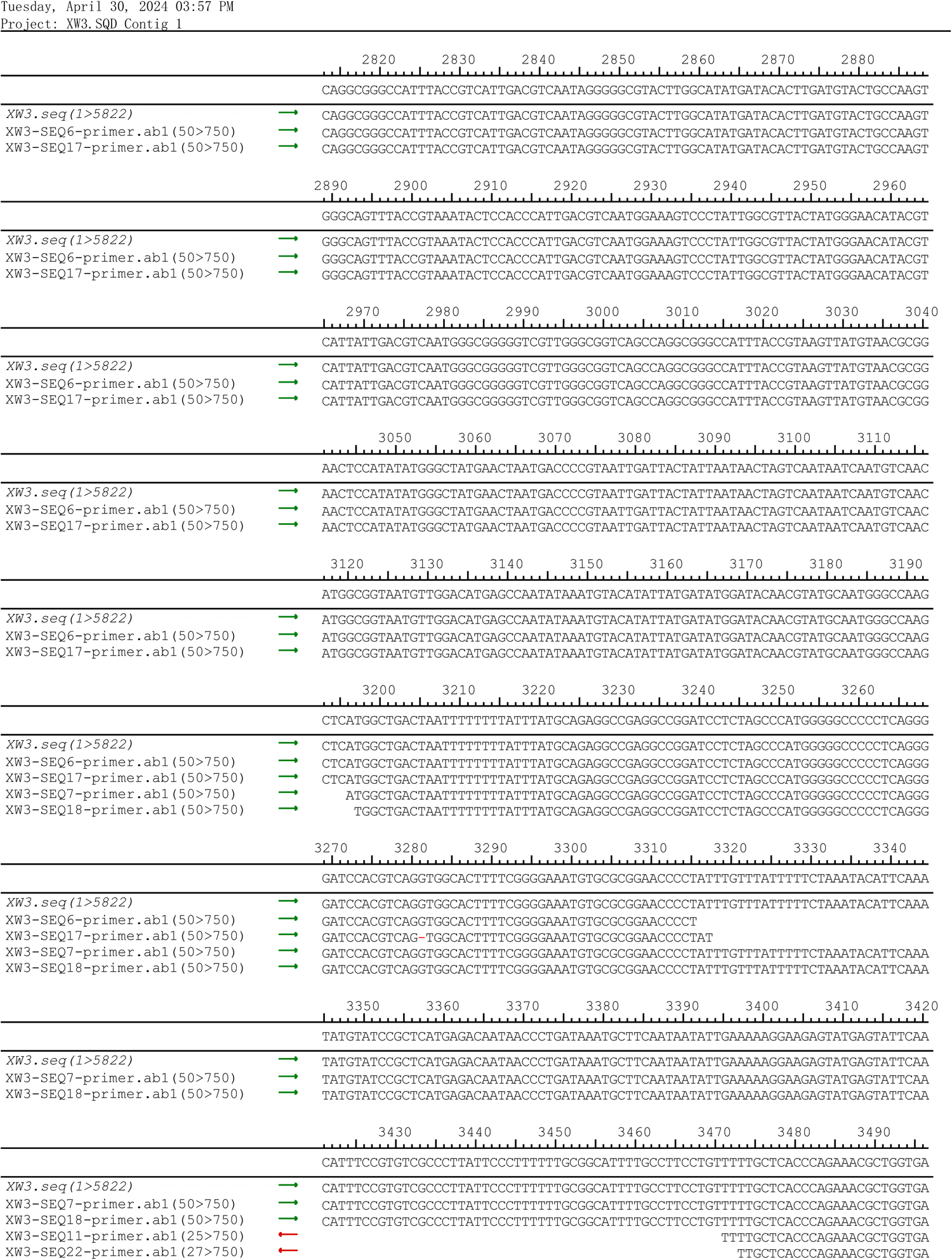

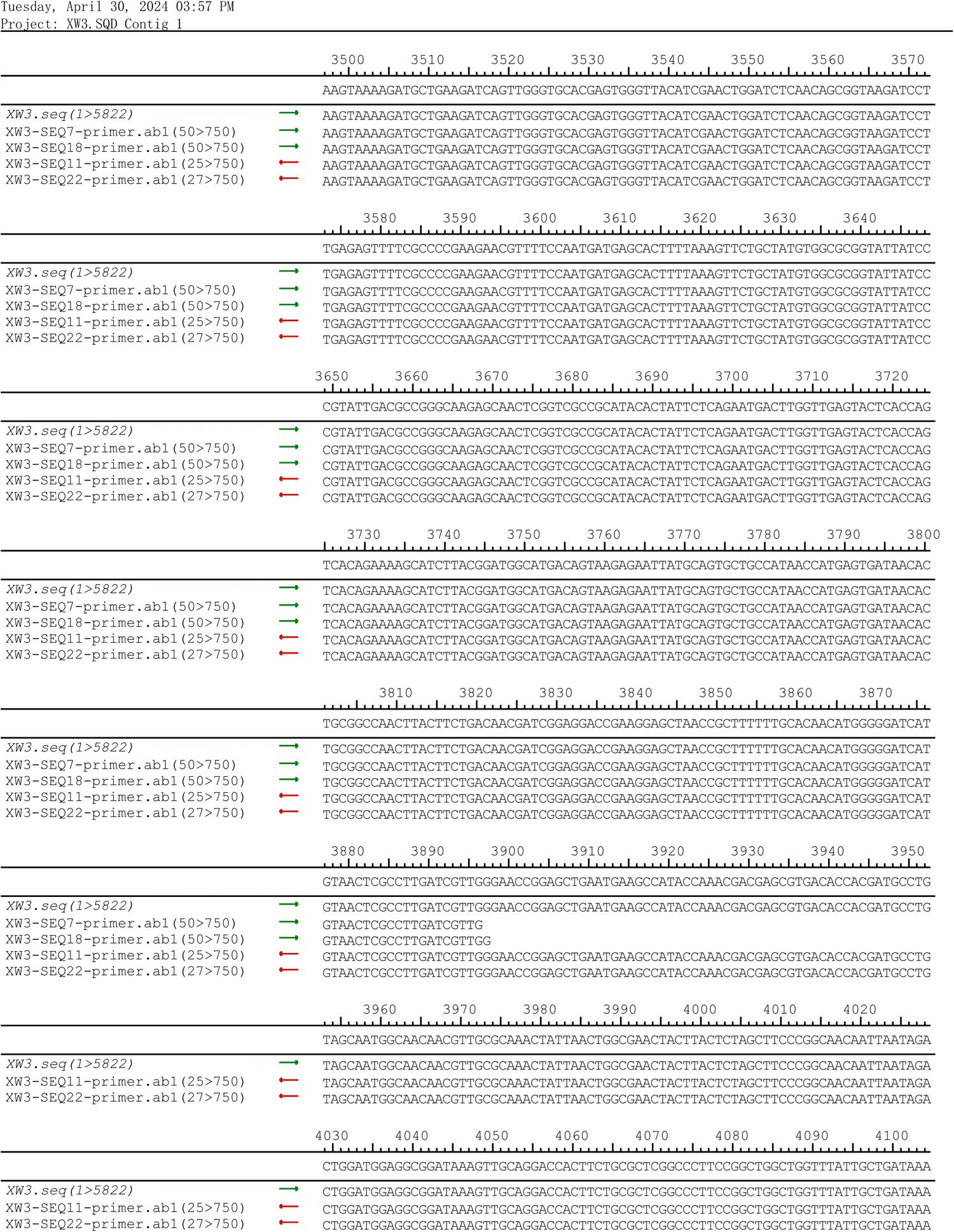

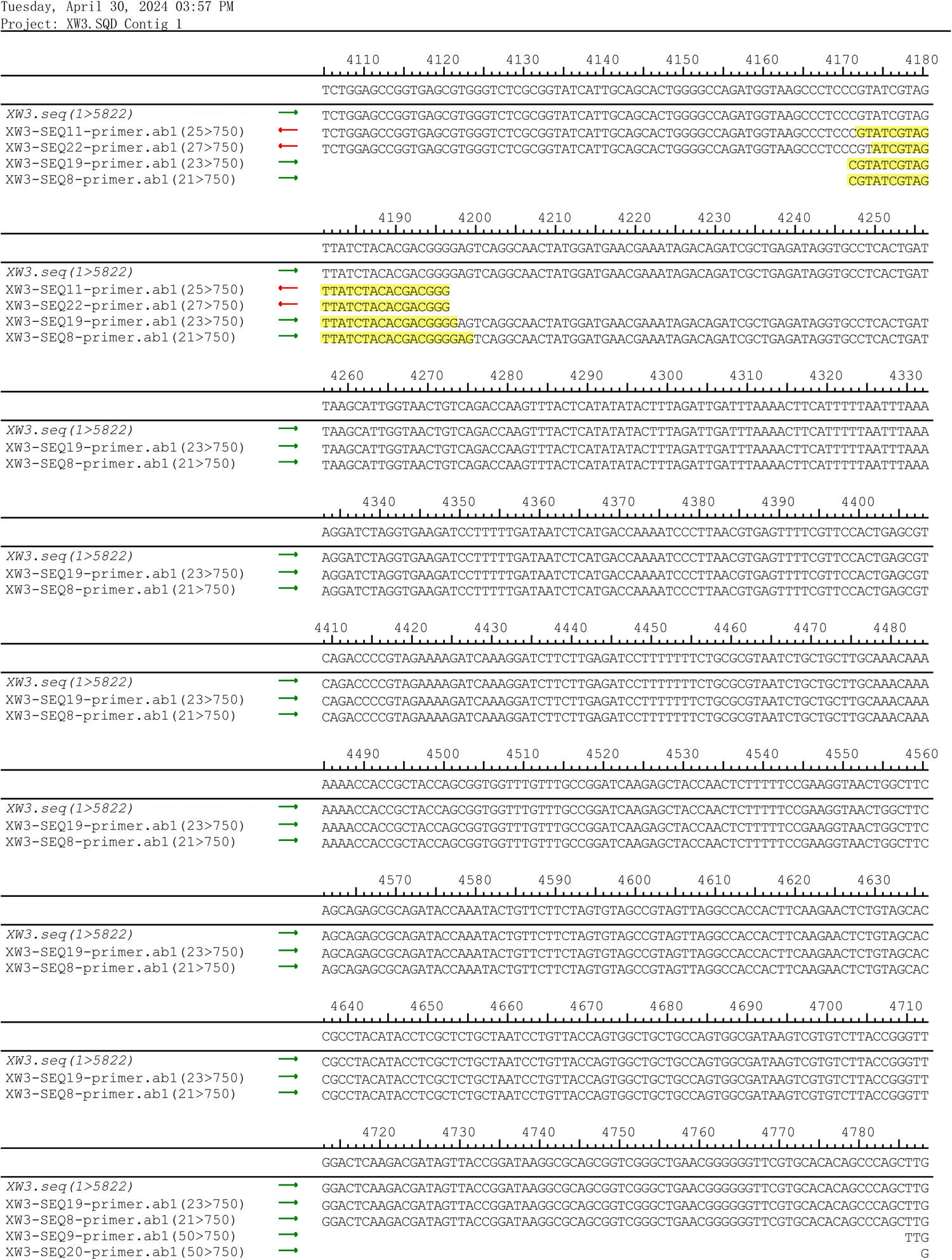

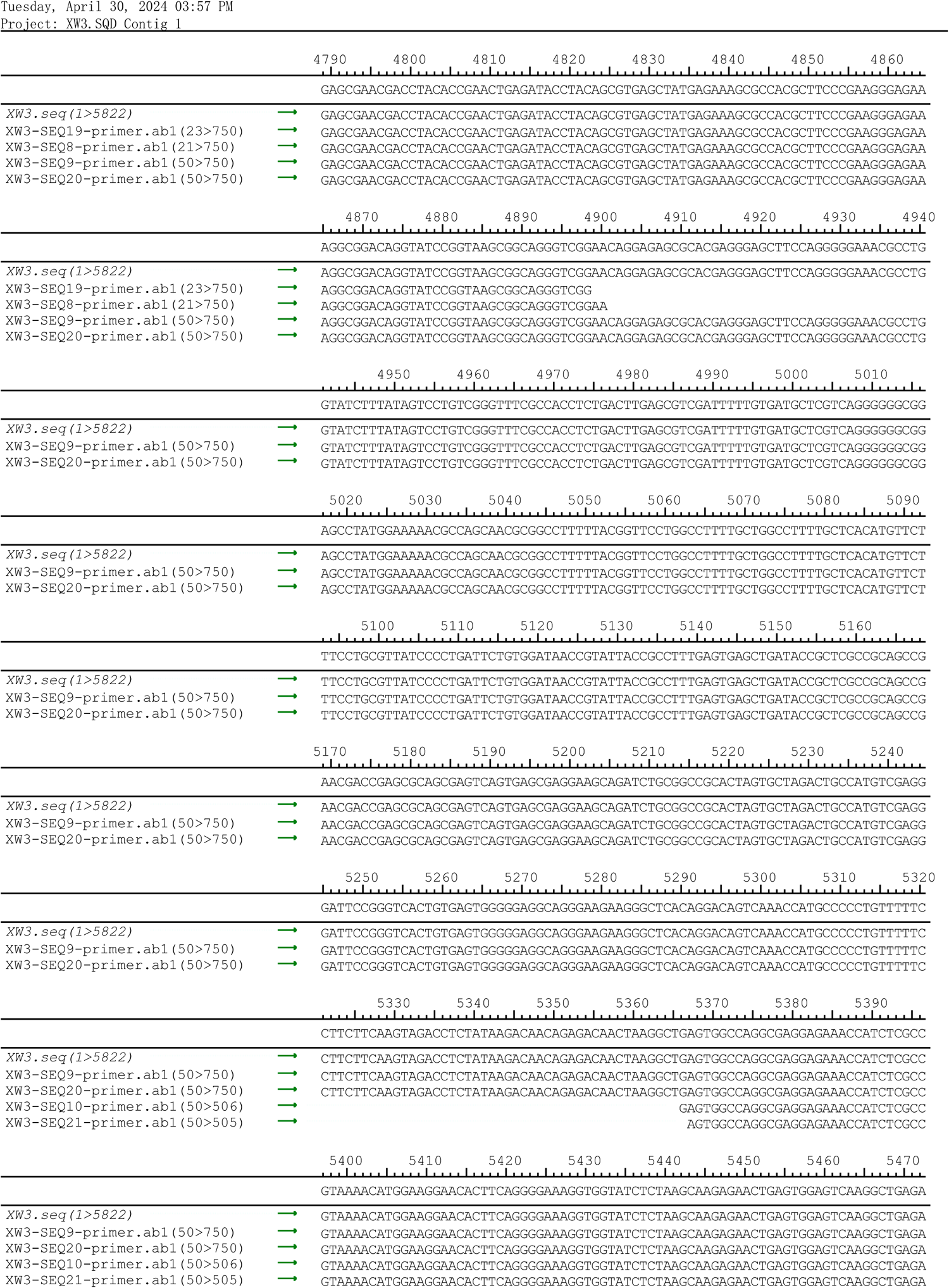

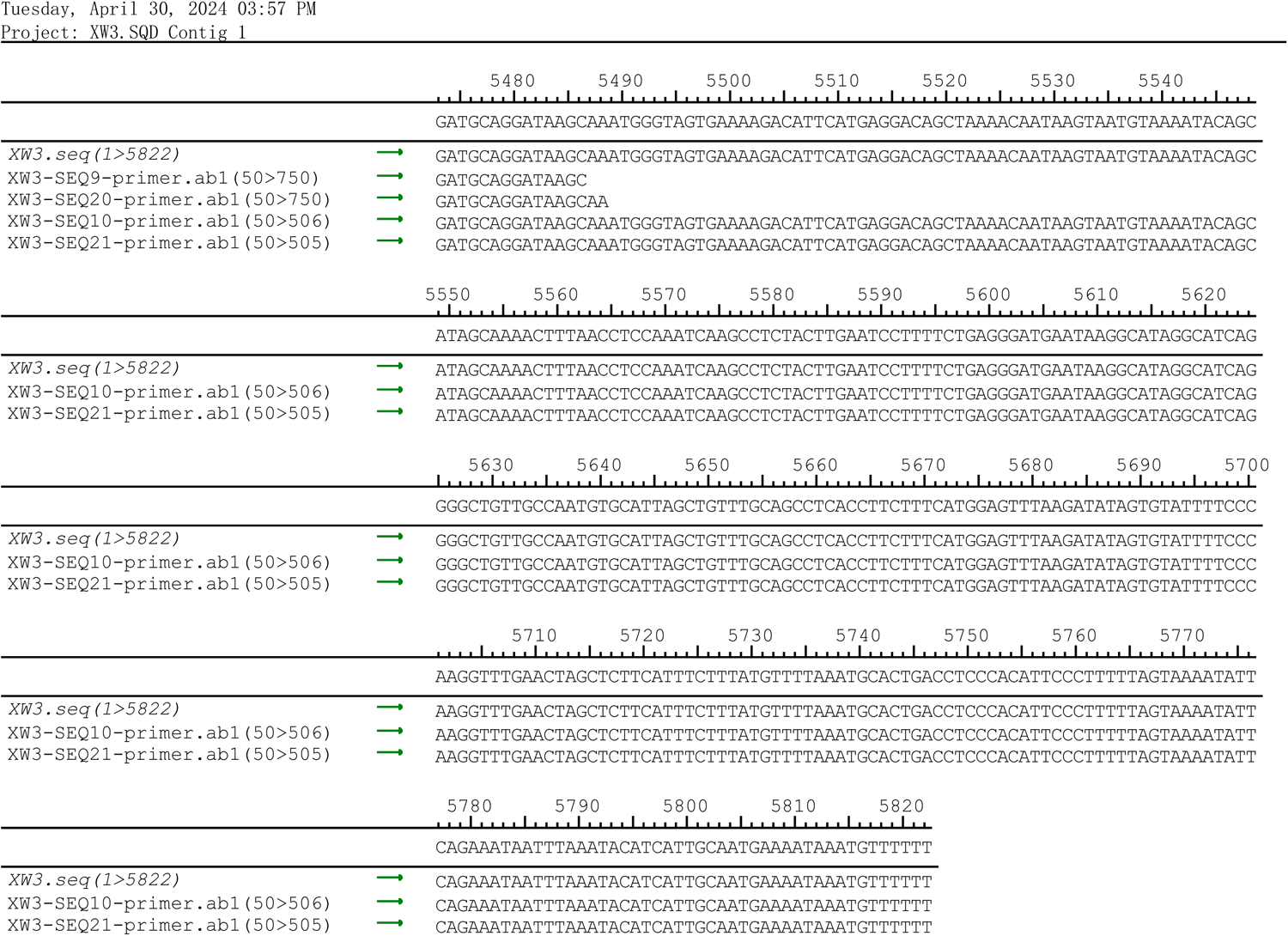

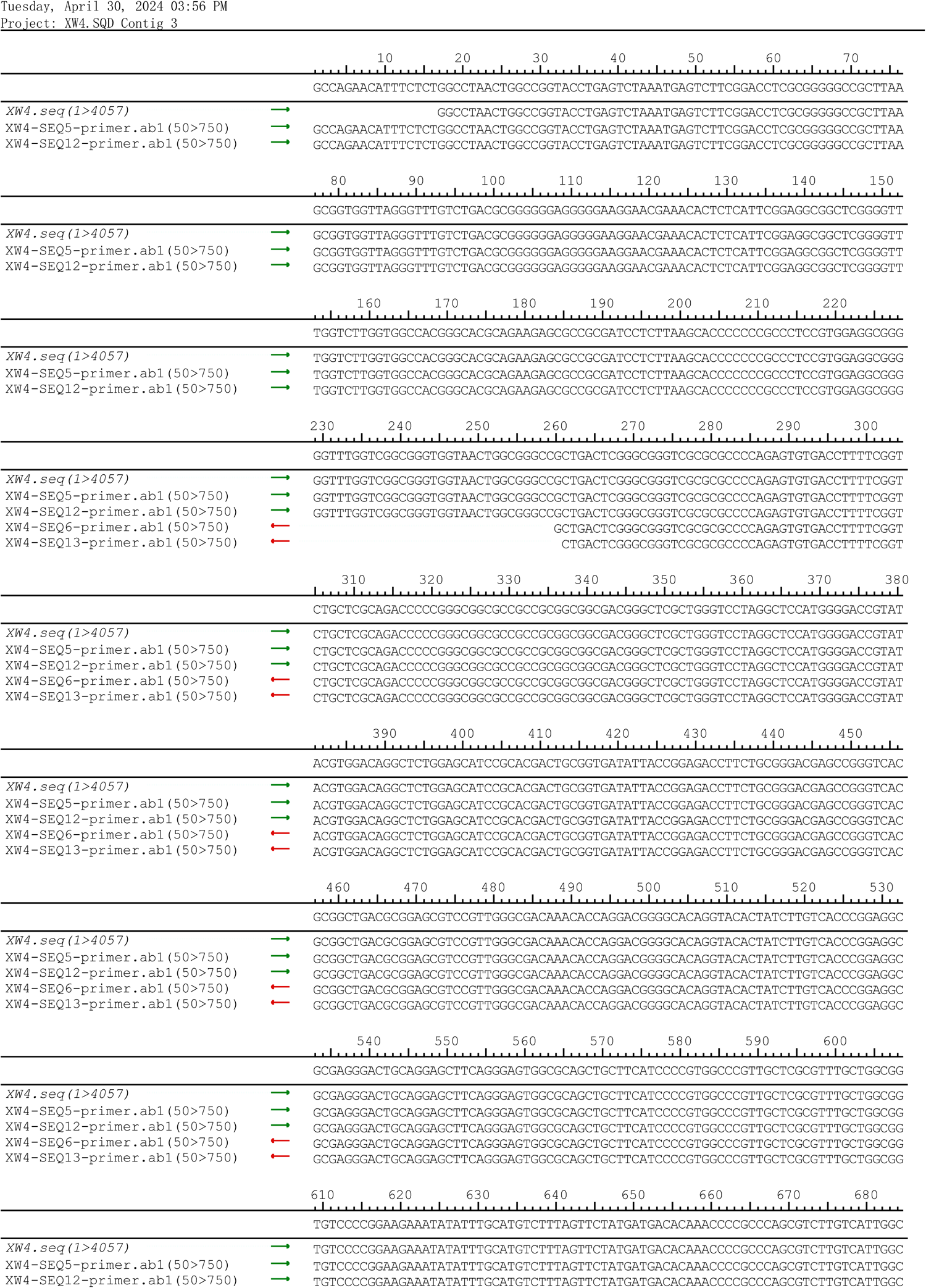

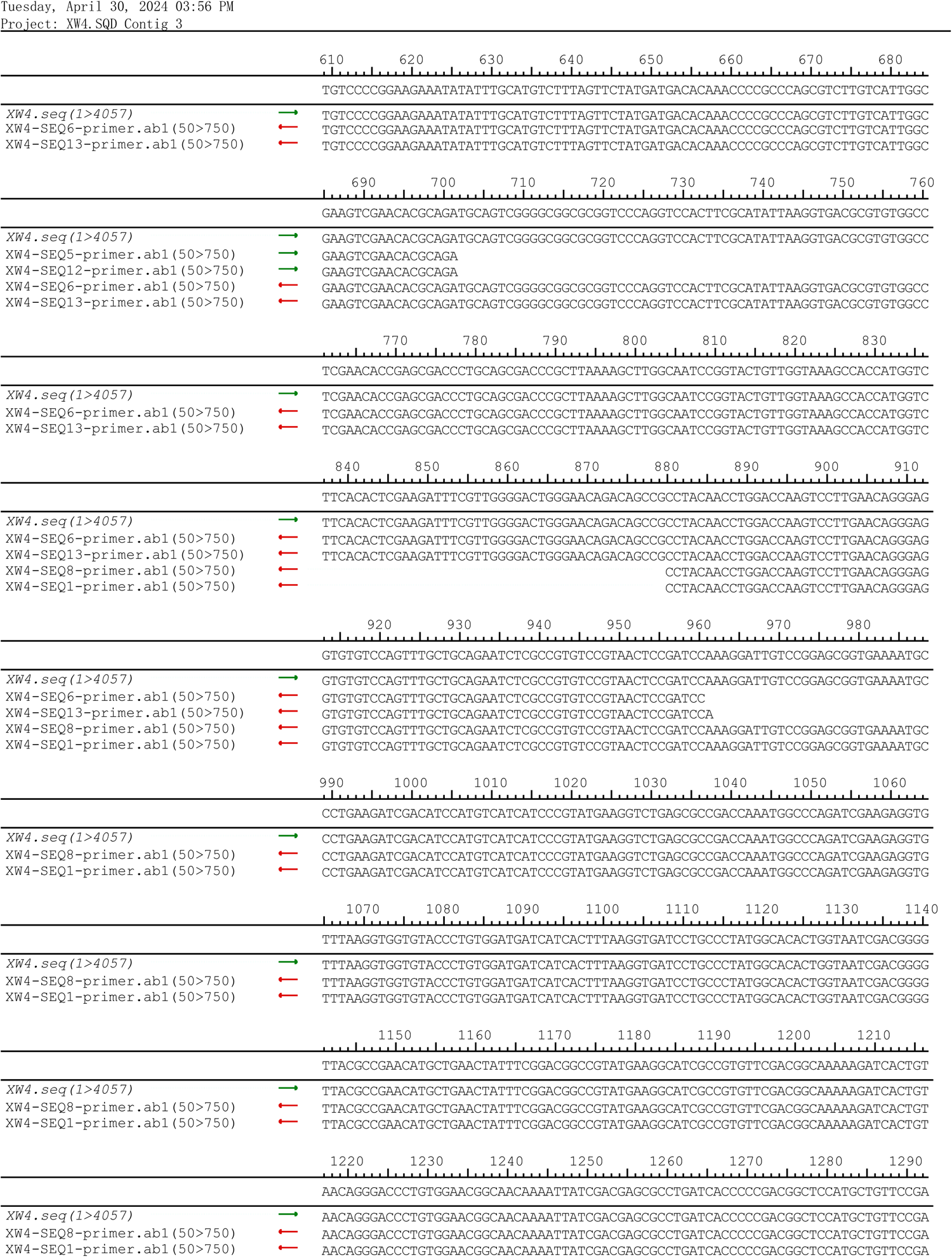

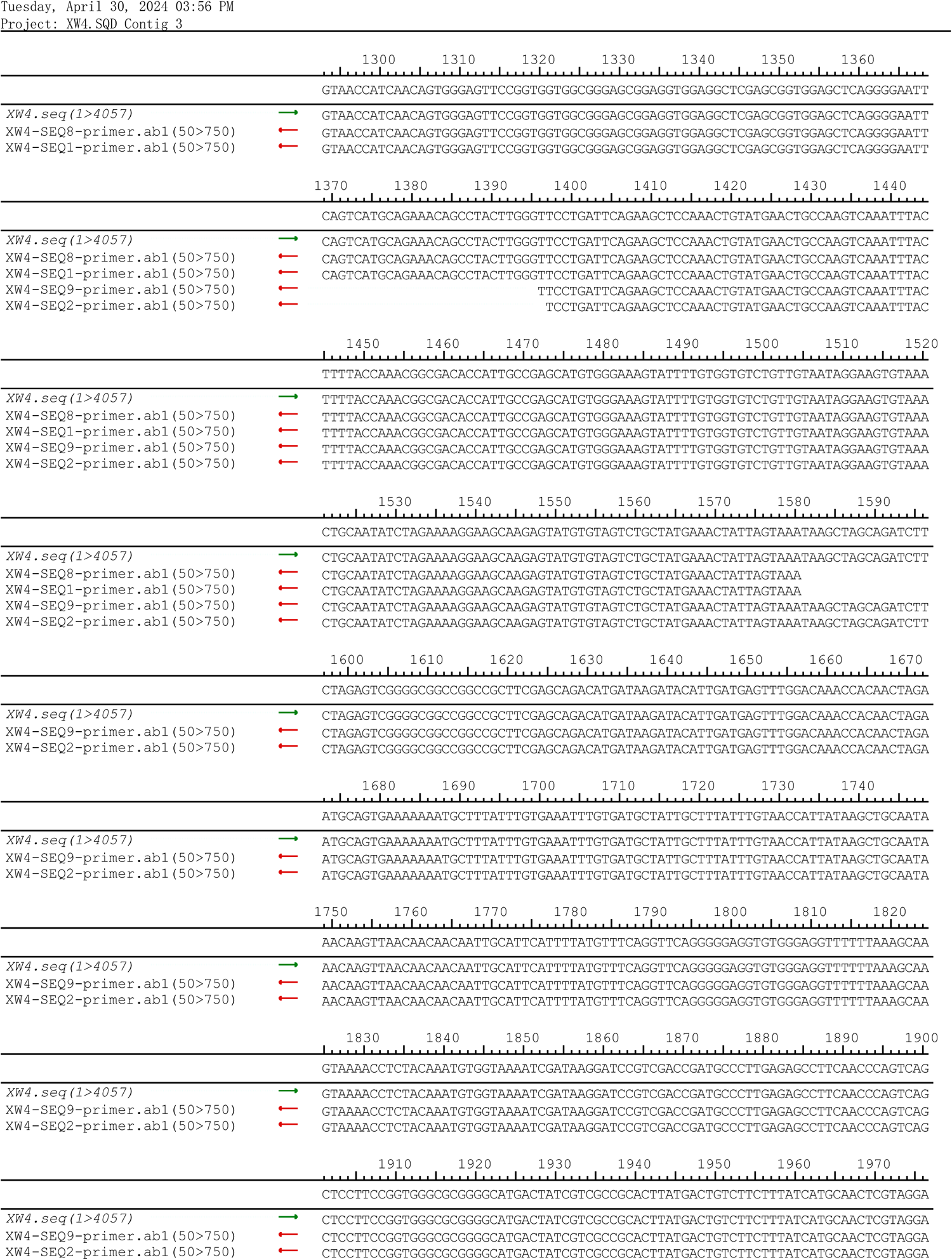

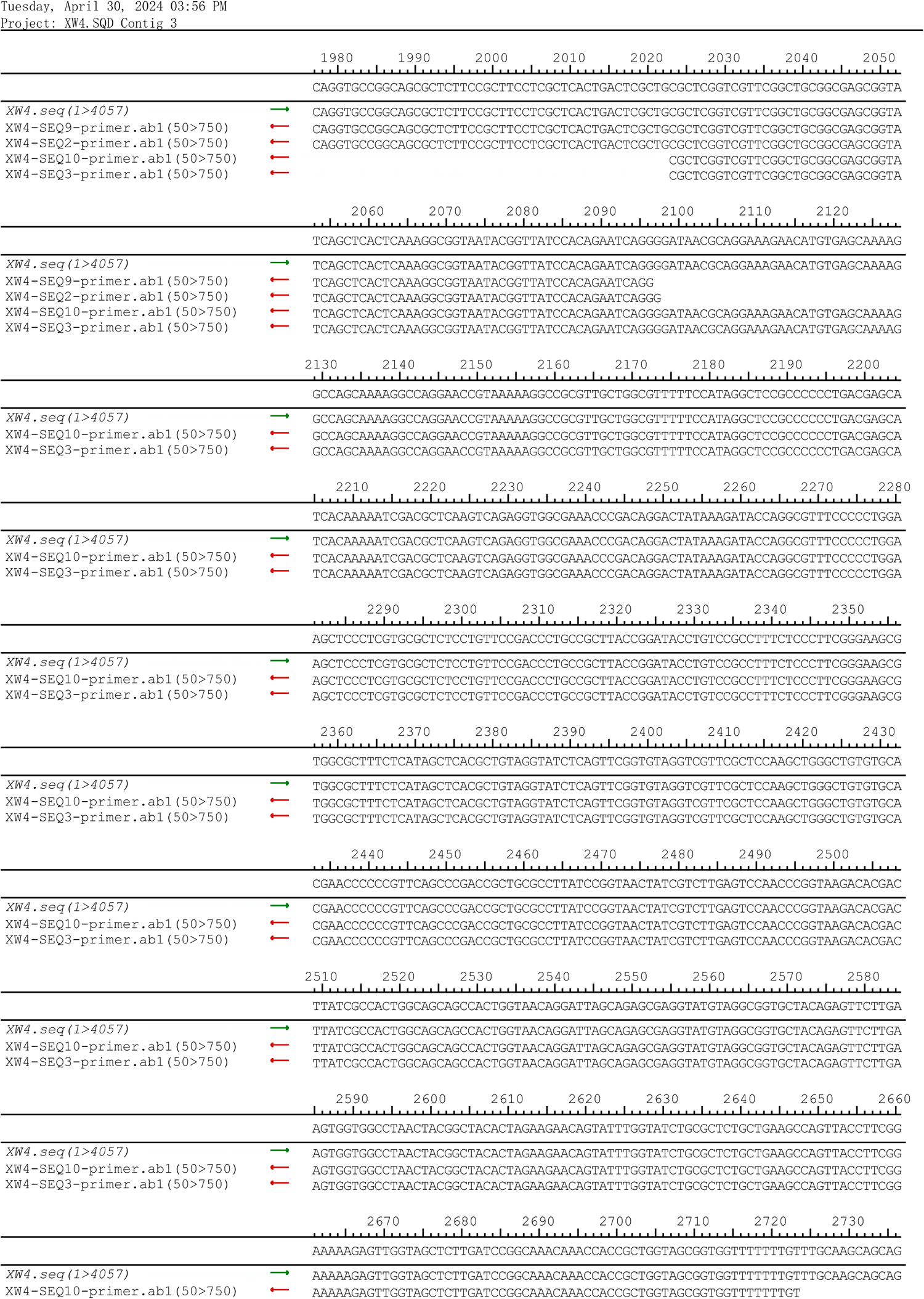

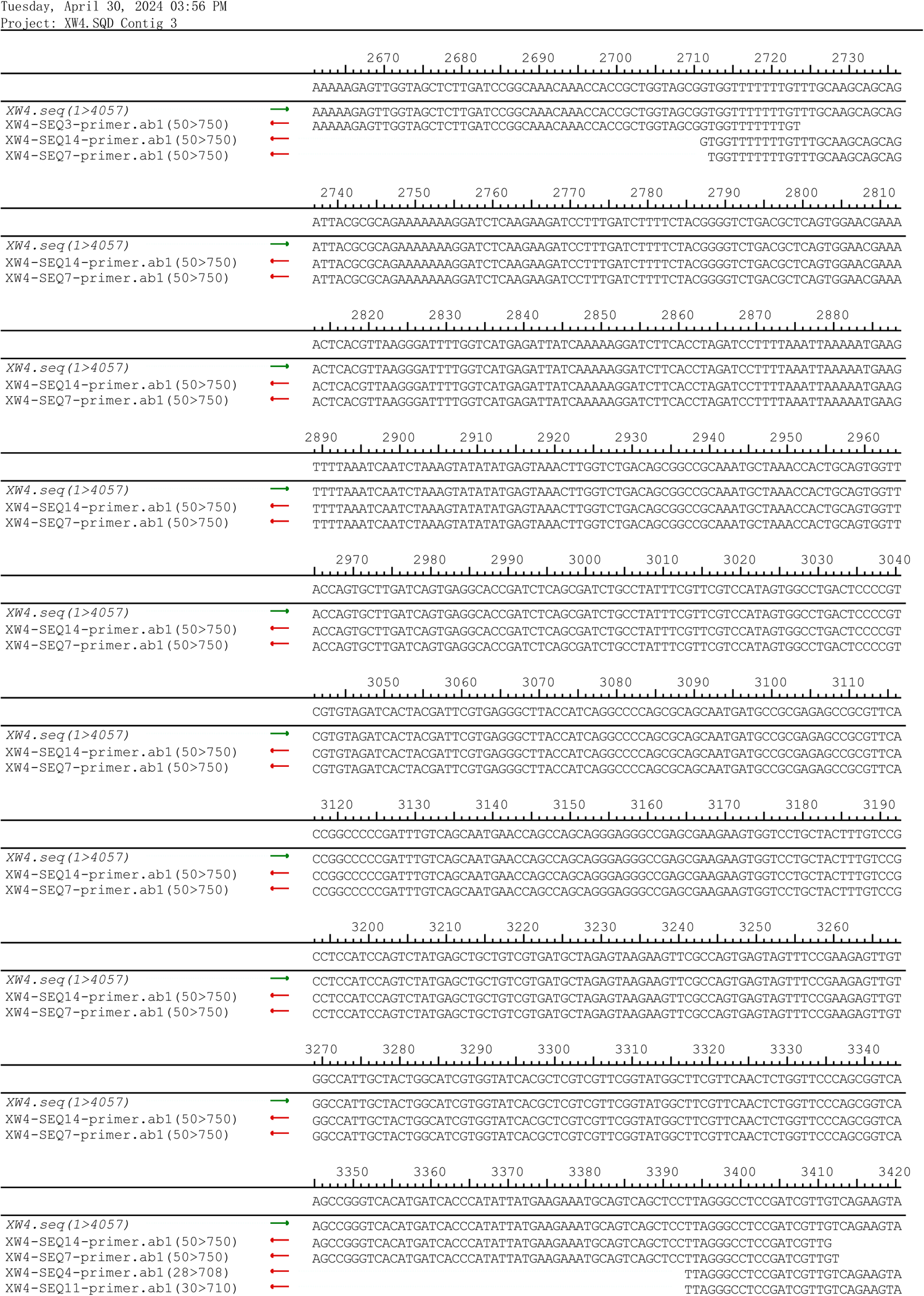

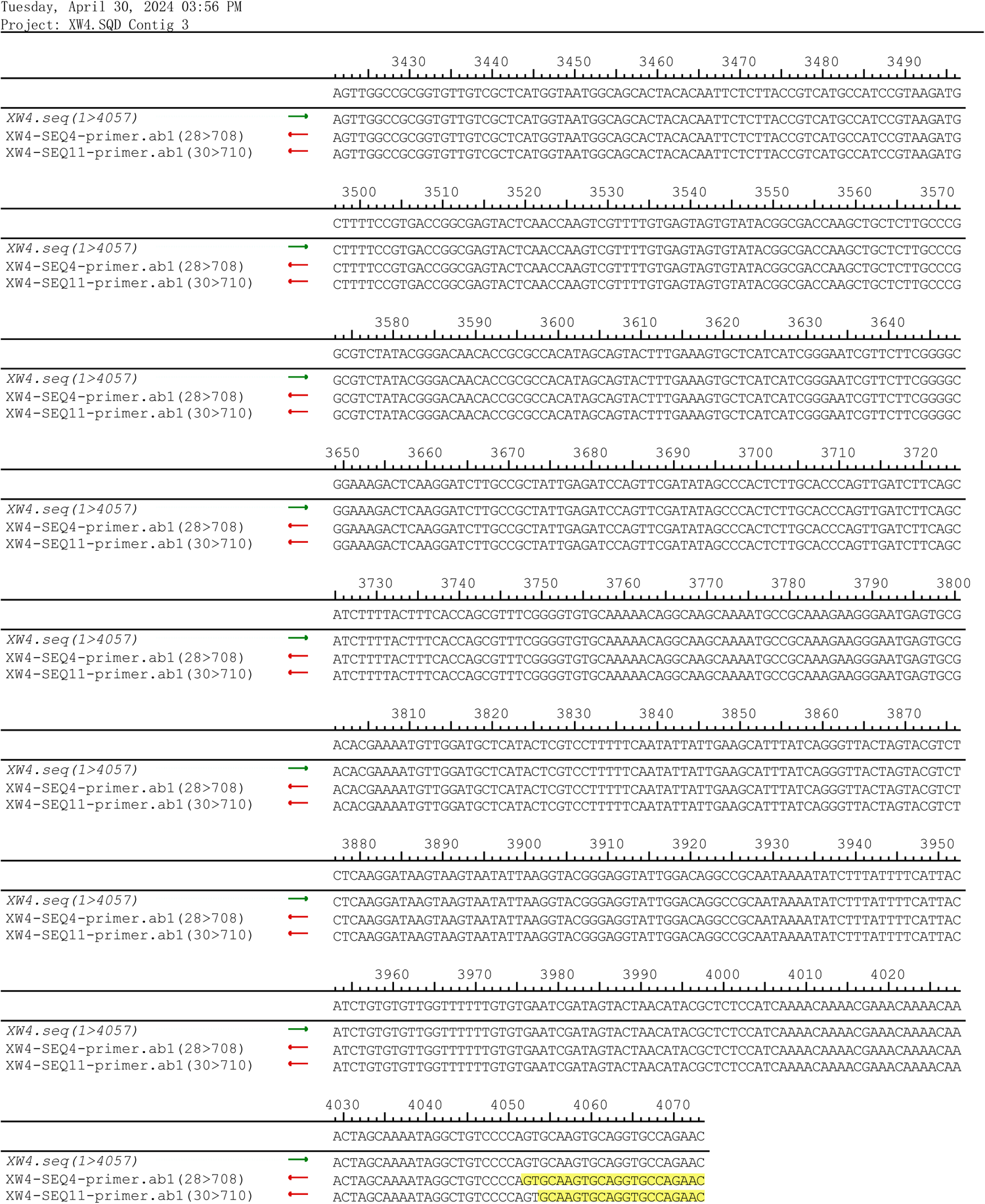

## Notes

### Competing Interest Statement

The authors have declared no competing interest.

